# *C9orf72* repeat expansion-carrying iPSC-microglia from FTD patients show increased phagocytic activity concomitantly with decreased number of autophagosomal-lysosomal vesicles

**DOI:** 10.1101/2024.08.14.607573

**Authors:** Hannah Rostalski, Tomi Hietanen, Dorit Hoffmann, Sami Heikkinen, Nadine Huber, Ashutosh Dhingra, Salvador Rodriguez-Nieto, Teemu Kuulasmaa, Sohvi Ohtonen, Henna Jäntti, Viivi Pekkala, Stina Leskelä, Petra Mäkinen, Kasper Katisko, Päivi Hartikainen, Šárka Lehtonen, Eino Solje, Jari Koistinaho, Tarja Malm, Anne M. Portaankorva, Teemu Natunen, Henna Martiskainen, Mari Takalo, Mikko Hiltunen, Annakaisa Haapasalo

**Affiliations:** A.I. Virtanen Institute for Molecular Sciences, University of Eastern Finland, Kuopio, Finland; Biotech Research & Innovation Center (BRIC), University of Copenhagen, Copenhagen, Denmark; Department of Pathology, Bartholin Institute, Copenhagen University Hospital, Copenhagen, Denmark (H.R.); Institute of Biomedicine, University of Eastern Finland, Kuopio, Finland; German Center for Neurodegenerative Diseases (DZNE), Tübingen, Germany; Institute of Clinical Medicine - Neurology, University of Eastern Finland, Kuopio, Finland; Neuro Center, Neurology, Kuopio University Hospital, Kuopio, Finland; Neuroscience Center, Helsinki Institute of Life Science (HiLIFE), University of Helsinki, Helsinki, Finland; Department of Neurology, University of Helsinki, Helsinki, Finland

**Keywords:** ALS, FTD, *C9orf72* hexanucleotide repeat expansion, microglia, phagocytosis, RNA sequencing

## Abstract

*C9orf72* hexanucleotide repeat expansion (HRE) is a major genetic cause of amyotrophic lateral sclerosis and frontotemporal dementia. The role of microglia in these *C9orf72* HRE-associated diseases is understudied. To elucidate effects of *C9orf72* HRE on microglia, we have characterized human induced pluripotent stem cell-derived microglia (iMG) from behavioral variant frontotemporal dementia (bvFTD) patients carrying the *C9orf72* HRE. *C9orf72* HRE iMG were compared to iMG from healthy controls and sporadic bvFTD patients. The phenotypes of iMG were analyzed using bulk RNA sequencing, biochemical and immunofluorescence analyses, and live cell imaging. *C9orf72* HRE-carrying iMG showed nuclear RNA foci and poly-GP dipeptide repeat proteins but no decreased C9orf72 mRNA or protein expression. TDP-43 pathology was absent from all bvFTD iMG. As compared to healthy control iMG, quantitative immunofluorescence analyses indicated that all bvFTD iMG had reduced number, size, and intensity of LAMP2-A-positive vesicles. *C9orf72* HRE-carrying iMG additionally showed decreased number, size, and intensity of p62/SQSTM1-positive vesicles. These changes were accompanied by increased phagocytic activity of the *C9orf72* HRE-carrying iMG. Serum starvation increased phagocytic activity also in the iMG of sporadic bvFTD patients. RNA sequencing revealed that iMG of *C9orf72* HRE-carrying bvFTD patients as compared to the iMG of sporadic bvFTD patients showed differential gene expression in pathways related to RNA and protein regulation and mitochondrial metabolism. Our data suggest potential alterations in the autophagosomal/lysosomal pathways in bvFTD patient iMG, which are further reinforced by the *C9orf72* HRE and functionally manifest as increased phagocytic activity.

## 1. Introduction

Microglia, the macrophages of the brain, perform key tasks during brain development, homeostasis, aging and disease. Their functions range from synaptic pruning and trophic support of neurons to immune functions, including pathogen defense, tissue repair, and cytokine signaling [1–4]. Since several risk genes for different neurodegenerative diseases are expressed by microglia [5], studying microglia-dependent disease mechanisms has gained great interest. During the last decade, protocols to generate human induced pluripotent stem cell (iPSC)-derived microglia (iMG) have been developed; see e.g., [6] for review. iMG-based model systems have already been used to study the impact of microglia on Alzheimer’s disease, Parkinson’s disease, frontotemporal dementia (FTD), amyotrophic lateral sclerosis (ALS), and Nasu-Hakola disease [7–13]. These models are important for elucidating the role of microglia in disease processes, since iMG express several genes associated with neurodegenerative diseases, including chromosome 9 open reading frame 72 (*C9orf72*), cluster of differentiation 33 (*CD33*), presenilin 2 (*PSEN2*), superoxide dismutase 1 (*SOD1*), TAR DNA binding protein *(TARDBP*), and triggering receptor expressed on myeloid cells 2 (*TREM2*) [14].

FTD and ALS are neurodegenerative diseases within the same neuropathological and genetic disease spectrum but with distinct clinical phenotypes. Whereas FTD predominantly affects the frontal and temporal regions of the brain and manifests by behavioral or linguistic changes followed by cognitive decline, patients with ALS suffer from degeneration of motor neurons [15–17]. Both, FTD and ALS can share an overlapping causative genetic background, including the hexanucleotide repeat expansion (HRE) in the *C9orf72* gene [18–20], and are often characterized by similar central nervous system (CNS) pathology, such as that involving accumulation of the TDP-43 protein [21, 22]. In a significant number of cases, FTD and ALS occur concomitantly and manifest as FTD-ALS phenotype [16]. Mechanistically, dysfunction in protein degradation pathways, including autophagy, has been suggested to lead to the accumulation of misfolded and toxic protein aggregates, underlying neuronal death and glial cell activation in FTD and ALS as well as other neurodegenerative diseases [23, 24]. This dysfunction could be linked to alterations in genes involved in autophagy initiation and vesicular trafficking, such as *C9orf72* [23–26].

Previous reports have indicated that *C9orf72* HRE-specific hallmarks, such as accumulation of RNA foci and dipeptide repeat (DPR) proteins and reduced *C9orf72* levels, indicating *C9orf72* haploinsufficiency, are present in neurons and, to some extent, glial cells [27–31]. Importantly, among CNS-resident cell types, the highest levels of the *C9orf72* gene are expressed in microglia [5, 32]. Microglial alterations have been linked to ALS and FTD [33], including increased microglia activation in the CNS areas undergoing *C9orf72* HRE-associated degeneration [34–36]. However, the role of microglia and the molecular mechanisms underlying their potential alterations in *C9orf72* HRE-associated ALS and FTD are still elusive. Microglia in *post-mortem* brains of *C9orf72* HRE carriers show an activated phenotype, enlarged lysosomes, and to some extent sense and antisense RNA foci [29, 32, 34]. Homo- and heterozygous knockout of the mouse *C9orf72* ortholog, *3110043O21Rik*, causes a severe peripheral autoinflammatory phenotype with inflammation and macrophage activation as well as hyper-reactivity to inflammatory stimuli. However, microglial activation was not consistently elevated as opposed to activation of macrophages in these mouse models [32, 37–40].

A recent report examining monocultures of iMG from ALS/FTD patients indicated moderate molecular and functional differences compared to healthy control iMG despite the presence of *C9orf72* HRE-associated pathologies [41]. On the other hand, iMG derived from *C9orf72* HRE carriers with ALS after stimulation with lipopolysaccharide [42] or AMPA [23] contributed to increased death of motoneurons. This was suggested to be related to impaired autophagy [23] and matrix metalloproteinase-9 secreted by the *C9orf72* HRE iMG [42]. These findings highlight the importance to study functions linked to or regulated by autophagy in the iMG of *C9orf72* HRE carriers. So far, to our knowledge, there are no studies on iMG derived from *C9orf72* HRE carriers with clinically diagnosed pure FTD.

In this study, our aim was to characterize the cell pathological and functional phenotypes of microglia from FTD patients. To this end, we have generated iMG from three healthy controls and three *C9orf72* HRE carriers and three non-carriers clinically diagnosed with behavioral variant frontotemporal dementia (bvFTD) for the first time. The iMG underwent detailed biochemical and cell biological analyses related to the *C9orf72* HRE-specific pathological hallmarks, TDP-43 pathology, RNA profiles, and microglial functionality. We found that iMG of the *C9orf72* HRE carriers showed the presence of RNA foci and DPR proteins but no signs of *C9orf72* haploinsufficiency. There was no apparent TDP-43 pathology in the iMG of *C9orf72* HRE-carrying or non-carrying bvFTD patients. Interestingly, the *C9orf72* HRE-carrying iMG showed changes in autophagosomal/lysosomal protein levels and significantly increased phagocytic activity. Global mRNA sequencing revealed differential expression of genes related to RNA, protein, and mitochondrial metabolism pathways in the iMG of *C9orf72* HRE bvFTD patients as compared to the iMG of sporadic bvFTD patients. Compared to iMG from healthy controls, iMG of *C9orf72* HRE-carrying bvFTD patients showed enrichment in genes related to acute inflammatory response. The present study, by comparing *C9orf72* HRE-associated and sporadic bvFTD iMG, provides completely new insights into differences in the gene expression and cellular function in these patient-derived immune cells.

## 2. Methods

### 2.1. Study subjects, skin biopsies, and *C9orf72* HRE genotyping

Skin biopsies were obtained at the Neurology outpatient clinic at the Neuro Center, Neurology, Kuopio University Hospital, Kuopio, Finland from FTD patients diagnosed with bvFTD clinical phenotype [17] and healthy control individuals. All donors gave a written informed consent. Fibroblast cultures were established from skin biopsies as previously described from three healthy individuals, three sporadic bvFTD patients not carrying the *C9orf72* HRE, and three *C9orf72* HRE-carrying bvFTD patients [43]. Repeat-primed PCR was performed on genomic DNA extracted from corresponding blood samples and skin biopsy-derived fibroblasts to confirm *C9orf72* HRE carriership (> 60 repeats) and non-carriers (< 30 repeats) [20]. Heterozygous *C9orf72* HRE carriership (> 145 repeats) was further confirmed in the iPSCs using AmplideX® PCR/CE C9orf72 Kit (49581; Asuragen). Since other bvFTD causative genetic factors are extremely rare among Finnish patients [44–46], samples were not tested for additional mutations. Collection of human monocytes and their differentiation to monocyte-derived microglia-like (MDMi) cells were performed according to Ryan KJ et al. [47].

### 2.2. Induced pluripotent stem cell (iPSC) generation and culturing

iPSCs were generated from fibroblasts as described previously [48] using CytoTune-iPS Sendai reprogramming kit (A16517, Thermo Fisher). In addition, one iPSC line of an FTD patient carrying the *C9orf72* HRE was purchased from EBiSC (cell line name: UCLi001-A, Biosamples ID: SAMEA3174431). The EBiSC Bank acknowledges University College London as the source of the human induced pluripotent cell line UCLi001-A, which was generated at UCL Queen Square Institute of Neurology with support from the NIHR UCLH Biomedical Research Centre, EFPIA companies and the European Union (IMI-JU). iPSCs were maintained on Corning® Matrigel® Basement Membrane Matrix Growth Factor Reduced (356230; Corning) coated 3.5 cm dishes (83.3900; Sarstedt) in E8 medium (Essential 8™ Medium [A1517001; Gibco™], supplemented with 0.5% [v/v] penicillin/streptomycin [15140122; Gibco™], 1×E8 supplement [A1517001; Gibco™]) at +37 L and 5% CO_2_. Cell culture medium was replaced once per day. Once confluency of 60-80% was reached, iPSCs were split by incubating for 3-5 min at +37 L in EDTA (15575020, Invitrogen) and kept in 5 µM Y-27632 2HCl (S1049, Selleck Chemicals) containing E8 medium. For freezing, iPSCs were collected in E8 medium supplemented with 10% (v/v) heat inactivated FBS (10500; Gibco™) and 10% (v/v) DMSO (D2650; Sigma) and stored long-term in liquid nitrogen. iPSCs were confirmed to be free of bacterial, fungal, or mycoplasma contaminations. To visualize nuclei, iPSCs were stained with 5 µM Vybrant™ DyeCycle™ Green Stain (V35004, Thermo Fisher) for 20 min at +37 L. Images using phase contrast and fluorescence at green channel (300 ms acquisition time) were taken using 10x objective (IncuCyte® S3; Essen BioScience). The iPSCs were confirmed by qPCR, immunofluorescence staining, and RNA sequencing to express pluripotency and stem cell markers (Supplementary Fig. 1-3). Detailed information on iPSCs is provided in Supplementary Table 1. Karyotype analysis by an experienced hospital geneticist indicated that two control iPSC lines (Con_1; Con_2) had non-pathological chromosomal alterations (in one, structurally abnormal short arm of chromosome 12; and in the other, short arm was replaced with long arm of chromosome 20, thereby forming an isochromosome 20q). The other iPSC lines showed normal karyotypes (Supplementary Fig. 4).

### 2.3. Differentiation of iPSCs into microglia and culturing of iMG

iPSCs were differentiated into iMG as previously described [13]. In brief, iPSCs were seeded as single cells and kept on Matrigel-coated dishes in Essential 8™ Medium (A1517001; Gibco™) supplemented with in 25 ng/ml Activin A (120-14E, Peprotech), 5 ng/ml BMP4 (120-05ET, Peprotech), 1 µM CHIR99021 (1386 B8, Axon), and 0.5% (v/v) penicillin/streptomycin (15140122; Gibco™), and in 5% O2 to induce mesodermal differentiation. The medium was supplemented with 10 µM (day 1) or 1 µM (day 2) Y-27632 2HCl (S1049, Selleck Chemicals), during the first 48 hours. Forty-eight hours after the initiation of differentiation, cells were kept in differentiation medium (Dulbecco’s Modified Eagle Medium:Nutrient Mixture F-12 [21331-020, Gibco™] containing 1x GlutaMAX [35050-038, Life Technologies], 0.5% [v/v] penicillin/streptomycin [15140-122, Gibco™], 64 mg/l ascorbic acid [A8960, Sigma-Aldrich], 14 µg/l sodium selenite [S5261, Sigma-Aldrich], 543 mg/l sodium bicarbonate [25080-094, Thermo Fisher]) and supplemented with 100 ng/ml FGF2 (AF-100-18B, Peprotech), 10 μM SB431542 (S1067, Selleckchem), 5 μg/ml insulin (I9278, Sigma-Aldrich), and 50 ng/ml VEGF (100-20A, Peprotech) and in 5% O_2_ to induce hemogenic differentiation. On days 4-7, cells were cultivated in differentiation medium supplemented with 50 ng/ml FGF2 (AF-100-18B, Peprotech), 5 μg/ml insulin (I9278, Sigma-Aldrich), 50 ng/ml VEGF (100-20A, Peprotech), 50 ng/ml TPO (300-18, Peprotech), 10 ng/ml SCF (300-07, Peprotech), 50 ng/ml IL-6 (200-06, Peprotech), 10 ng/ml IL-3 (200-03, Peprotech), and under 18% O_2_. On day 8, erythromyeloid progenitors were transferred to ultra-low attachment dishes (4615, Corning®). Medium (IMDM [21980032; Gibco™] supplemented with 10% [v/v] heat-inactivated FBS [10500; Gibco™], 0.5% [v/v] penicillin/streptomycin [15140122; Gibco™], 5 μg/ml insulin [I9278, Sigma-Aldrich], 5 ng/ml MCSF [300-25, Peprotech], 100 ng/ml IL-34 [200-34, Peprotech]) was changed every other day and cell numbers were kept under 3.5 million cells per dish. On day 16, iMG were seeded in microglia medium (IMDM [21980032; Gibco™] supplemented with 10% [v/v] heat-inactivated FBS [10500; Gibco™], 0.5% [v/v] penicillin/streptomycin [15140122; Gibco™], 10 ng/ml MCSF [300-25, Peprotech], 10 ng/ml IL-34 [200-34, Peprotech]) on PDL (P6407; Sigma) coated vessels for phagocytosis experiments (96 well plates [167008; Thermo Fisher]), fluorescence *in situ* hybridization (FISH; 13 mm cover glasses [631-1578; VWR] in 24 well plates [150628; Thermo Fisher]), immunofluorescence studies (µ-Slide 8 Well [80826; Ibidi], 13 mm cover glasses in 24 well plates, or 9 mm cover glasses [631-0169; VWR] in 48 well plates [150687; Thermo Fisher]), or RNA and protein extraction (18 mm cover glasses [631-0153; VWR] in 12 well plates [142475; Thermo Fisher]). For protein extraction and FISH, erythromyeloid progenitors were frozen on day 8, 10, or 12 and stored in liquid nitrogen in freezing medium (10% [v/v] DMSO in microglia medium). After thawing, the differentiation protocol was continued starting from the same day of the protocol on which the cells were frozen (Supplementary Fig. 5).

### 2.4. Protein extraction and Western blotting

iMG were seeded on day 16 (100,000 cells/well) and harvested on day 22. For experiments with Bafilomycin A1 treatment, iMG were treated on day 22 with 150 nM Bafilomycin A1 (B1793-10UG, Sigma Aldrich) for 6 h at +37 L before harvesting. Cells were washed twice with ice-cold DPBS (17-512F; Lonza), scraped in lysis buffer (10 mM Tris–HCl, 2 mM EDTA, 1% [m/v] sodium dodecyl sulphate [SDS]; 1:100 protease and 1:100 phosphatase inhibitors [1862209 and 1862495; Thermo Scientific]). Samples were stored at −20 L. Thawed samples were sonicated (2 cycles [10 s], 30 s between cycles, high setting; Bioruptor Next Gen, Diagenode) and heated (+90 °C for 7 min). Protein concentrations were measured using bicinchoninic acid assay (23225; Thermo Scientific) and plate reader (Infinite M200; Tecan Group Ltd.). Samples (approx. 15 µg/lane) and molecular weight marker (26616; Thermo Scientific) were supplied with 20% (v*/*v) 2-mercaptoethanol and 1×NuPAGE LDS sample buffer (NP0007; Invitrogen), heated for 7 min at +90 °C, and loaded on 4-12% Bis-Tris gels (NP0335; Invitrogen) for sodium dodecyl sulfate-polyacrylamide gel electrophoresis (80 V for 15 min, 110 V for 1 h 15 min). Proteins were transferred on 0.2 μm polyvinylidene fluoride membranes (1704157; Bio-Rad) using Trans-Blot Turbo Transfer System (Bio-Rad; 25 V, 1.0 A, 30 min). Blots were blocked either in 5% (w*/*v) milk powder or 5% (w*/*v) BSA (A9647; Sigma-Aldrich) diluted in 1×TBST (139.97 mM NaCl, 24.94 mM Tris, 0.5% (v*/*v) Tween 20; pH 7.4) for 1 h at room temperature (RT) according to the recommendations of primary antibody manufacturers. Blots were incubated overnight at +4 °C with the following primary antibodies diluted in 1×TBST: anti-C9orf72 (1:500, GTX634482; GeneTex), anti-LAMP2-A (1:1,000, ab18528; Abcam), anti-LC3B (1:3,000, ab51520; Abcam), anti-phospho-TDP-43 (1:3,000, TIP-PTD-P02; CosmoBio), anti-SQSTM1 (1:1,000, 5114S; CST), anti-TDP-43 (1:2,000, 10782-2-AP; Proteintech). Next, blots were incubated with species-specific horseradish peroxidase-linked secondary antibodies (1:5,000, NA935, NA934V or NA931V; GE Healthcare) for 1 h at RT, and for 5 min with suitable enhanced chemiluminescence substrate solutions (RPN2236 or RPN2235; GE Healthcare; UltraScence Pico Ultra [CCH345; BioHelix]). ChemiDoc MP Imaging System (Bio-Rad) was used to detect the chemiluminescence signals. For each sample, intensity values per area of bands of interest were quantified using Image Lab^TM^ (6.0.0; Bio-Rad). Values were normalized to total protein signals above approx. 18 kDa in size from blots stained with No-Stain™ Protein Labeling Reagent (A44449, Invitrogen). Data are shown as % of Con_2 line (mean value of each experiment set to 100%).

### 2.5. Fixation, FISH, immunofluorescence staining, and confocal microscopy

Cells were fixed on day 22 in 4% (v/v) formaldehyde (28908; Thermo Scientific) in DPBS for 20 min at +37 L, washed two times with DPBS, and stored at +4 L. FISH was performed similarly as described previously [49] using a hybridization temperature of +55 L and fluorescently labeled locked nucleic acid TYE^TM^ 563-(CCCCGG)_3_ probe to detect the sense foci. TYE^TM^ 563-(CAG)_6_ probe was used as a negative control probe to verify the specificity of the TYE^TM^ 563-(CCCCGG)_3_ probe against the *C9orf72* HRE sequence (Exiqon). H_2_O added to samples for FISH was pre-treated with diethyl pyrocarbonate (D5758; Sigma).

For immunofluorescence staining, fixed cells were permeabilized using 0.1% (v/v) Triton X-100 in DPBS for 10 min at RT on a rocker and washed three times with DPBS. For blocking, cells were incubated in blocking solution (1.5% or 5% [for C9orf72] [v/v] goat immunoglobulin G isotype control [02-6202; Invitrogen] or 1% [for p62/SQSTM1] or 5% [for TREM2] [w/v] BSA in DPBS) for 30 min at RT on a rocker. Anti-C9orf72 was diluted in 1% (w/v) BSA in DPBS. All other primary and secondary antibodies were diluted in blocking solution and incubated with the cells either at RT for 1.5 h or overnight at +4 L followed by washing three times with DPBS. The following primary antibodies were used: anti-C9orf72 (1:500, GTX634482; GeneTex), anti-CX3CR1 (1:500, 1H14L7; Thermo Fisher), anti-IBA1 (1:250, MA5-27726; Thermo Fisher), anti-LAMP2-A (1:200, ab18528; Abcam), anti-NANOG (1:100, AF1997; R&D Systems), anti-P2RY12 (1:250, HPA014518; Sigma-Aldrich), anti*-*POU5F1 (1:400, MAB4401; EMD Millipore), anti-SSEA4 (1:400, MAB4304; EMD Millipore), anti-SQSTM1 (1:200, sc-28359; Santa Cruz Biotechnology), anti-TDP-43 (1:500, 10782-2-AP; Proteintech), anti-TMEM119 (1:250, HPA051870; Thermo Fisher), anti-TRA-1-81 (1:200, MAB4381; EMD Millipore), anti-TREM2 (1:50, PA5-46980; Thermo Fisher). The following fluorescently labeled species-specific secondary antibodies were used at 1:300 or 1:500 dilution: Alexa Fluor^®^ 488 (A-11001, A-11006, A-11034, A-11055, and A-11059; Invitrogen), Alexa Fluor® 568 (A-11004, A-11057; Invitrogen), Alexa Fluor^®^ 647 (A-21244; Invitrogen). To stain the cell nuclei, cells were mounted in 4′,6-diamidino-2-phenylindole (DAPI)-containing mounting medium (H-1800; Vector Laboratories) or incubated in 0.2 µg/ml DAPI solution (D952, Sigma) in DPBS for 5 min at RT on rocker. For C9orf72, LAMP2-A, p62/SQSTM1 and TDP-43 staining, cells were mounted in phalloidin-containing mounting medium (H-1600; Vector Laboratories). Next, 8-bit images with a resolution of 15.2841 pixels per micron were acquired using laser scanning confocal microscopy (Plan-Apochromat 63x/1.40 Oil DIC M27 objective, Axio Observer.Z1/7, LSM800; Zeiss), or 16-bit images with a resolution of 6.2016 pixels per micron using EC Plan-Neofluar 40x/1.30 Oil DIC M27 objective and Axio Observer.Z1 microscope (Zeiss). Same microscopy settings were used for all samples (including negative control samples where no primary antibody was used) and replication experiments.

### 2.6. Image analysis

Microglia were stained for TDP-43, C9orf72, p62/SQSTM1, or LAMP2-A and stained with phalloidin and DAPI as described above. Confocal microscopy images were processed using Fiji (2.3.0/1.53f51; [50]). Phalloidin images were used to create masks, which outlined the cell bodies and allowed measurement of the total cell body area per image. For this, auto local thresholding (version 1.10, Sauvola method, radius = 15, default *k* and *r* values) was applied, followed by Gaussian Blur filter (sigma = 2 [sigma = 4 for C9orf72 images]) and removal of particles with a radius smaller than 5 µm. To fill holes within cell bodies, particles of a size between 0-4 µm were included. After watershed segmentation, regions of interest were outlined without further area or shape exclusion. DAPI images were used to indicate nuclei and measure nuclear areas per image (for TDP-43 analysis). To segment nuclei, background subtraction (rolling ball radius =100 pixels), Gaussian Blur filter (sigma = 2), auto thresholding (Renyi’s entropy method), and removal of particles with a radius smaller than 5 µm were applied followed by fill holes command.

#### 2.6.1. C9orf72 intensity analysis

C9orf72 signal was quantified as sum intensities of the fluorescence signal originating from the fluorescently labeled secondary antibody within phalloidin-positive areas per image. Sum intensities were normalized to phalloidin-positive area per image. Values from samples without primary antibody incubation (negative control) were subtracted from values from samples with primary antibody incubation.

#### 2.6.2. TDP-43 translocation analysis

TDP-43 signals were quantified as sum intensities of the fluorescence signal originating from the secondary antibodies in nuclear (DAPI-positive) and cytosolic areas (nuclear signal subtracted from signal within whole cell body [phalloidin-positive areas]) for each image. Sum intensities were normalized to nuclear and cytosolic area sizes, respectively for each image. To determine unspecific signal intensities for each cell line, intensity values for nucleus and cytosol were obtained from samples stained without primary antibody (negative control). Maximum unspecific signal intensities for each cell line per experiment were calculated. Cytosolic to nuclear TDP-43 signal ratio was calculated by subtracting the maximum unspecific area-normalized sum intensities for each line from nuclear and cytosol signals, respectively. To determine the extent of TDP-43 translocation into the cytosol, a recently developed TDP-43 translocation analysis was used [51]. In brief, in this analysis the signal of nuclear and cytosolic TDP-43 is categorized into four stages (none, mild, moderate, severe TDP-43 translocation from nucleus to cytosol; Supplementary Fig. 6 a). The same concept was used here to categorize TDP-43 signals. For this, the number of translocation-positive cells was manually counted on the basis of nuclear and cytosolic TDP-43 intensity values. For the manual analysis, TDP-43 stages “none” and “mild” were grouped as “non-pathological” and “moderate” and “severe” as “pathological”. This was applied because the cytosolic to nuclear TDP-43 intensity signals of manually selected cells did not differ between “none” and “mild” nor “moderate” and “severe” groups (Supplementary Fig. 6 b) but differed between the groups “non-pathological” and “pathological” (Supplementary Fig. 6 c).

#### 2.6.3. p62/SQSTM1 and LAMP2-A particle analysis

Signals were quantified as sum intensities of the fluorescence signal originating from the secondary antibody normalized to p62/SQSTM1- or LAMP2-A-positive particle areas. Mean size of p62/SQSTM1- or LAMP2-A positive particles and mean number of particles per total cell body area within phalloidin-positive areas per image were calculated. Background subtraction (rolling ball radius =10 pixels) and thresholding (Maximum Entropy algorithm [p62/SQSTM1]; Renyi’s entropy method [LAMP2-A]) were applied so that a minimal number of particles were detected in samples stained without primary antibody (negative control). Only particles with a minimum area of 0.5 µm^2^ were included in the analysis (p62/SQSTM1).

### 2.7. Phagocytosis assay

On day 16, iMG were plated on 96-well plates (approx. 30,000 cells/well). For serum starvation, 24 h prior to assay, iMG were kept in serum-free microglia medium. On day 23 or 24, cells were incubated in serum-free microglia medium supplemented with zymosan-coupled pH-sensitive fluorescent particles (P35364, Invitrogen) at final concentrations of 62.5 µg/ml. Cells without particles were used for background subtraction in the red channel. Microscopy images were acquired with 20x objective from brightfield, and red fluorescent channels (300 ms acquisition time) every 30 min using IncuCyte® S3 (Essen BioScience). Per experiment, four images per well per biological replicate (3-4) per line were acquired. For analysis, the IncuCyte® software (version 2019B) was used. For bright field channel, segment adjustment of 0.5 was used. For the red channel, Top-Hat segmentation (radius 10 µm, 2 RCU threshold) and edge sensitivity of −60 were used. To estimate the phagocytic activity, the size of the red fluorescent area within cells or intensity values of red channel (Total Red Integrated Intensity [RCU x µm²/image]) were calculated and normalized to total cell area (before beads were added) for each image. Values were calculated for each biological replicate and pooled from two independent differentiations.

### 2.8. DPR immunoassay

iMG were seeded on day 16 (400,000-500,000 cells/well) and snap frozen on dry ice after removing medium on day 23. Sample preparation and the DPR immunoassay were carried out on the Meso Scale Discovery (MSD) platform as described by previously [52]. Total protein levels in all samples were used to normalize the MSD assay signals. In total, one control iMG line (Con_3) and two sporadic iMG lines (Ftd_1, Ftd_4) as *C9orf72* HRE non-carrying controls, and all three *C9orf72* HRE carrier iMG lines (Ftd_5, Ftd_6, Ftd_7), were included in the immunoassay.

### 2.9. RNA extraction

iPSCs were detached with EDTA, centrifuged for 10 min at 8,000 x g, resuspended, and processed in NucleoProtect RNA (740400.250, MACHEREY-NAGEL) as described by the manufacturer, and stored at −80 L until RNA extraction. On day 16, iMG were plated (225,000 cells/well). On day 23, iMG were washed twice with ice-cold DPBS and scraped in 200 µl ice-cold DPBS. RNA extraction was conducted directly. Total RNA was isolated (11828665001, Roche Molecular Systems, Inc.), and RNA concentrations were measured using NanoDrop^TM^ One (Thermo Scientific).

### 2.10. RNA sequencing and data processing

Bulk RNA sequencing (RNA-seq) was performed using RNA extracted as described above from iPSCs and iMG. Library preparation and RNA sequencing was conducted by Novogene (UK) Company Limited as described previously [53]. Sequencing yielded 19.9–32.4 million sequenced fragments per sample.

The 150 nt pair-end RNA-seq reads were quality controlled using FastQC (version 0.11.7) (https://www.bioinformatics.babraham.ac.uk/projects/fastqc/). Reads were then trimmed with Trimmomatic (version 0.39) [54] to remove Illumina sequencing adapters and poor quality read ends, using as essential settings: ILLUMINACLIP:2:30:10:2:true, SLIDINGWINDOW:4:10, LEADING:3, TRAILING:3, MINLEN:50. Reads aligning to mitochondrial DNA or ribosomal RNA, phiX174 genome, or composed of a single nucleotide, were removed using STAR (version 2.7.9a) [55]. The remaining reads were aligned to the Gencode human transcriptome version 38 (for genome version hg38) using STAR (version 2.7.9a) [55] with essential non-default settings: --seedSearchStartLmax 12, --alignSJoverhangMin 15, --outFilterMultimapNma× 100, --outFilterMismatchNmax 33, -- outFilterMatchNminOverLread 0, --outFilterScoreMinOverLread 0.3, and --outFilterType BySJout. The unstranded, uniquely mapping, gene-wise counts for primary alignments produced by STAR were collected in R (version 4.1.0) using Rsubread::featureCounts (version 2.8.1) [56], ranging from 14.7 to 24.3 million per sample. After normalization, using varianceStabilizingTransformation (from DESeq2 version 1.32.0), the data were subjected to sample-level quality control and a minor batch effect was identified between sequencing batches (Supplementary Fig. 7). Batch-corrected DEGs between experimental groups were identified in R (version 4.1.2) using DESeq2 (version 1.32.0) [57] by employing Wald statistic and lfcShrink for FC shrinkage (type=“apeglm”) [58]. Pathway enrichment analysis was performed on the gene lists ranked by the pairwise DEG test log2FCs using command line GSEA (GSEA version 4.1.0) [59] with Molecular Signatures Database gene sets (MSigDB, version 7.5.1) [29].

### 2.11. Data processing, visualization, and statistical analyses

To test whether data points within experimental groups were normally distributed, Shapiro-Wilk test was used. Bartlett’s test was used to test for equality of variances among groups. To test for significance between two different experimental groups, two-tailed independent samples t-test (for normally distributed data and equal variances) or Mann-Whitney U test was performed. To test for significance between more than two groups, one-way analysis of variance (ANOVA) followed by Tukey’s multiple comparisons test (variance stabilized counts) or Sidak’s multiple comparisons test (phagocytosis assay) was used if data points were normally distributed and variances equal between groups. Otherwise Kruskal-Wallis test followed by Dunn’s multiple comparisons test was used. Two-way ANOVA followed by Sidak’s multiple comparisons test was used for testing statistical significance related to treatment effect among the experimental groups (Bafilomycin A1 treatment). To assess statistical independence between categorical variables, Chi-Square test was used. All tests were performed using GraphPad Prism software (version 8.4.3 for Windows, GraphPad Software, San Diego, California USA, www.graphpad.com). Test statistics (*F* value for ANOVA; *H* value for Kruskal-Wallis test; and corresponding *p*-values) are reported in the figure legends. P-values for pairwise comparisons are indicated above the compared groups within the figures. P*-*values < 0.05 were considered significant. Graphs were created using the GraphPad Prism software (version 8.4.3 for Windows). Number of independent differentiations of iMG from iPSCs are indicated in each figure legend. Total number of statistical units per group is indicated as “*n*”. Statistical units are depicted as individual data points. Microscopy images shown in the manuscript were processed using ImageJ (1.53f51).

### 2.12. Ethics statement

The patient data used in this project were pseudonymized and handled using code numbers only as indicated in Supplementary Table 1. Donors of skin biopsies gave their written informed consent prior to sample collection. The study was performed in accordance with the principles of the declaration of Helsinki. Obtaining skin biopsies, establishment of the biopsy-derived fibroblast cultures, and generation of iPSCs were performed with the permission 123/2016 from the Research Ethics Committee of the Northern Savo Hospital District, Kuopio, Finland. The authors declare no competing interests.

## 3. Results

### 3.1. iPSC-derived microglia of bvFTD patients and healthy controls acquire microglial identity

In this study, we used a previously published protocol [13] to generate microglia (iMG) from iPSCs of three healthy donors, three bvFTD patients carrying the *C9orf72* HRE, and three bvFTD patients not carrying the *C9orf72* HRE (sporadic). The study subject characteristics are listed in Supplementary Table 1. The iPSCs showed typical colony formation and expression of pluripotent stem cell marker genes (Supplementary Fig. 1-3), whereas these markers were not expressed in the differentiated iMG anymore (Supplementary Fig. 3). After 23 days of differentiation, iMG expressed markers, which have been suggested to be unique to human primary microglia [60] as opposed to CNS-associated macrophages [61], and used in other studies to confirm myeloid lineage/microglial identity [10, 13, 14, 62, 63] on RNA level (Fig. 1 a). Expression of a subset of microglial genes on protein level, i.e., CX3CR1, IBA1, P2RY12, TMEM119, and TREM2, was confirmed using immunofluorescence staining after 22 days in culture (Fig. 1 b, Supplementary Fig. 8). According to visual inspection by microscopy, 100% of iMG cells expressed these markers. Furthermore, global mRNA sequencing analysis indicated that the iMG clustered close to MDMi cells but separately from iPSCs or monocytes, from which the iMG or MDMi were differentiated, respectively (Fig. 1 c). Altogether, these characterizations confirmed that the generated iMG harbored microglial identity based on their high and specific expression of well-established microglial markers.

**Fig. 1.**
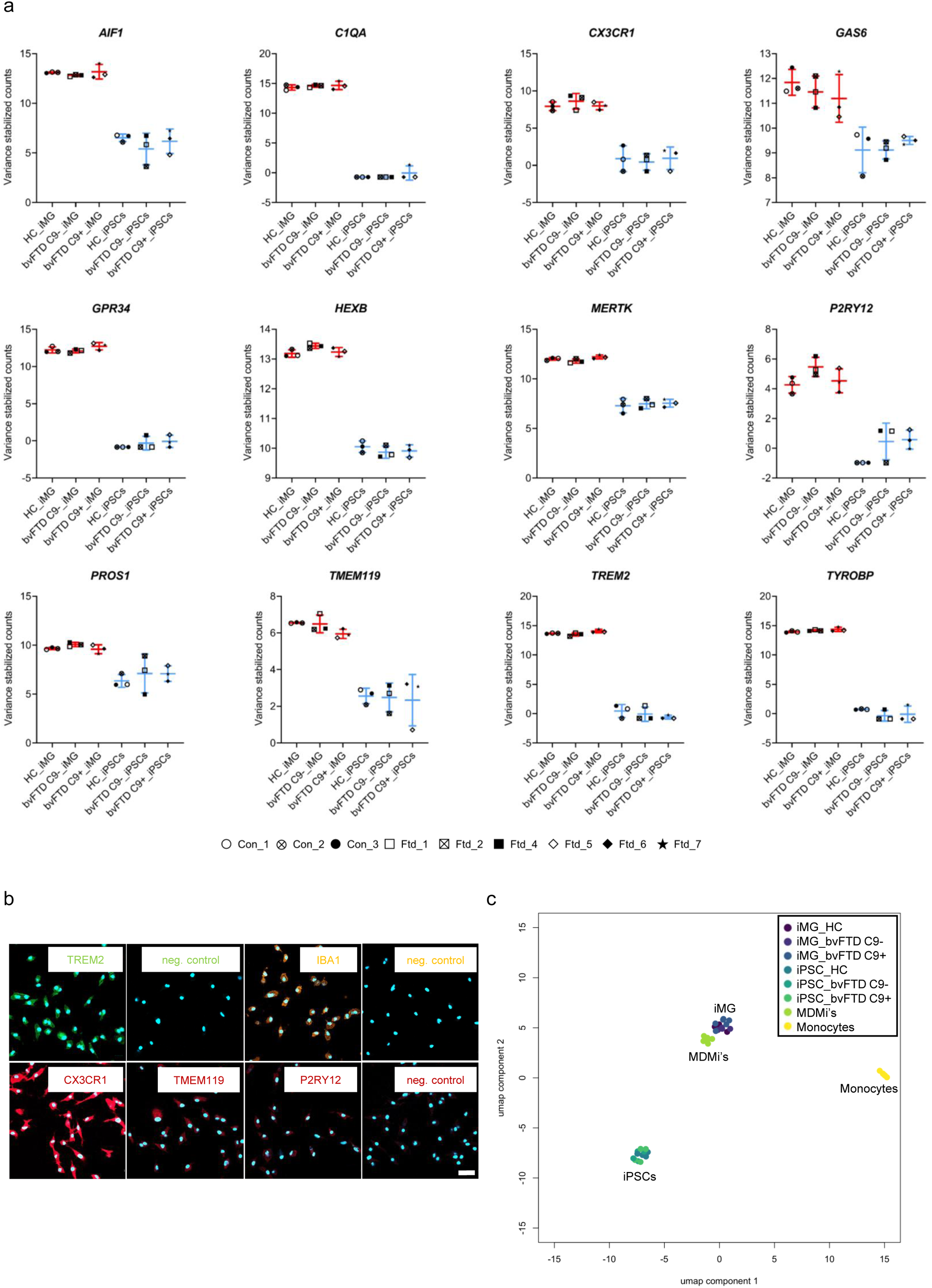
Microglial identity of iMG from healthy controls, sporadic (C9-) and *C9orf72* HRE-carrying (C9+) bvFTD patients. (a) RNA sequencing data for each donor, displayed as variance stabilized counts, show expression of selected microglial genes in iMG (red) but not in the iPSCs (blue). (b) iMG of a healthy donor (Con_1) were stained for specific microglial marker proteins. The corresponding negative control stainings (no primary antibody) are shown as indicated. Scale bar = 40 µm. Please see Supplementary Figure 8 for the staining of these markers in the iMG from the other donors. (c) UMAP plot shows clustering of iMG closely to monocyte-derived microglial cells (MDMi) but separately from monocytes and iPSCs. Data are shown from one differentiation batch. The individual iPSC and iMG lines are indicated by different symbols below the plots in (a). Abbrev.: *AIF1* = Allograft inflammatory factor 1; *C1QA* = Complement C1q subcomponent subunit A; bvFTD = behavioral variant frontotemporal dementia; CX3CR1 = CX3C chemokine receptor 1; *GAS6* = Growth arrest – specific 6; *GPR34* = G-protein coupled receptor 34; *HEXB* = Beta-hexosaminidase subunit beta; HRE = hexanucleotide repeat expansion; IBA1 = ionized calcium binding adapter molecule 1; iMG = induced pluripotent stem cell-derived microglia; iPSC = induced pluripotent stem cell; MDMi = monocyte-derived microglia; *MERTK* = MER Proto-Oncogene, Tyrosine Kinase; P2RY12 = P2Y purinoceptor 12; *PROS1* = Protein S; TMEM119 = transmembrane protein 119; TREM2 = Triggering receptor expressed on myeloid cells 2; *TYROBP* = TYRO protein tyrosine kinase-binding protein; UMAP = Uniform Manifold Approximation and Projection

### 3.2. iMG of *C9orf72* HRE carriers do not show reduced C9orf72 levels

*C9orf72* HRE has been shown to result in *C9orf72* haploinsufficiency in the patients, indicated by the reduction of C9orf72 mRNA expression by approximately 50% in the frontal cortex and lymphoblast cells of HRE carriers [18]. Also, concurrently reduced C9orf72 RNA and protein levels in the CNS of *C9orf72* HRE carriers have been reported in some studies [64, 65]. Microglia express the highest levels of *C9orf72* compared to other brain cells [32], and *C9orf72* has been suggested to be involved in key cellular functions that are executed by microglia, such as phagocytosis [66]. Hence, we investigated whether the iMG of *C9orf72* HRE carriers showed reduced total C9orf72 mRNA or protein levels. RNA sequencing showed that the iMG of the *C9orf72* HRE carriers did not harbor reduced *C9orf72* RNA levels as compared to the iMG of the non-carrier healthy controls or sporadic bvFTD patients (Fig. 2 a). In line with these RNA-level findings, Western blot analysis did not indicate significant differences in the C9orf72 long protein isoform levels (54 kDa) between the *C9orf72* HRE-carrying *vs.* non-carrying iMG (Fig. 2 b, c). Moreover, immunofluorescence staining of C9orf72 did not show significant differences in the levels of C9orf72 in the iMG of *C9orf72* HRE carriers as compared to the non-carriers either (Fig. 2 d, e). Altogether, these results suggest that *C9orf72* haploinsufficiency is not detected at the RNA or protein level in the iMG of *C9orf72* HRE carriers.

**Fig. 2.**
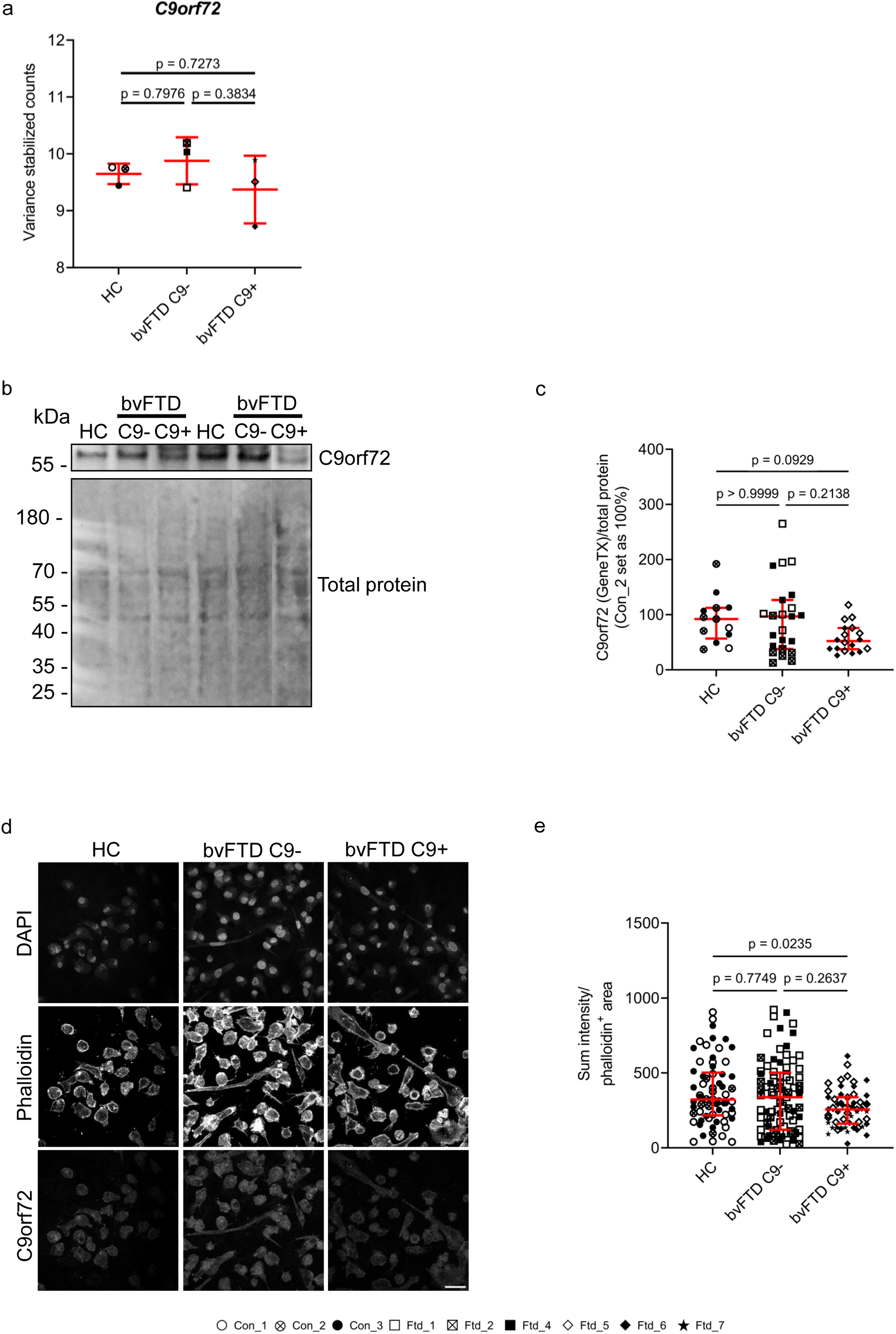
C9orf72 levels in iMG from healthy controls, sporadic (C9-) and *C9orf72* HRE-carrying (C9+) bvFTD patients. (a) *C9orf72* gene expression for each donor according to RNA sequencing shown as variance stabilized counts; *F* = 1.031, *p* = 0.4121; *n* [HC] = 3; *n* [bvFTD C9-] = 3; *n* [bvFTD C9+] = 3. (b) A representative Western blot image of C9orf72 protein levels in iMG. (c) For quantification of Western blot images, C9orf72 levels were normalized to total protein levels and to mean of normalized C9orf72 levels of samples from a healthy control. Each data point represents one biological replicate from three independent differentiation batches in total; *H* = 5.405, *p* = 0.0670; *n* [HC] = 13; *n* [bvFTD C9-] = 23; *n* [bvFTD C9+] = 18. (d) Representative immunofluorescence images of C9orf72. Scale bar = 30 µm. Phalloidin signal was used to normalize the quantified C9orf72 levels to cell confluency per region. (e) Quantification of C9orf72 levels based on immunofluorescence images. Each data point represents mean C9orf72 levels per region from three independent differentiation batches; *H* = 6.208, *p* = 0.0449; *n* [HC] = 65; *n* [bvFTD C9-] = 81; *n* [bvFTD C9+] = 57. For each group, descriptive statistics are shown as mean ± standard deviation (a) or median ± interquartile range (c). The individual iMG lines are indicated by different symbols at the bottom. One-way ANOVA followed by Tukey’s multiple comparisons test (a) or Kruskal-Wallis test followed by Dunn’s comparisons test (c, e) were used for statistical analysis. Abbrev.: bvFTD = behavioral variant frontotemporal dementia; DAPI = 4′,6-diamidino-2-phenylindole; HC = healthy control; HRE = hexanucleotide repeat expansion; iMG = induced pluripotent stem cell-derived microglia

### 3.3. iMG of C9orf72 HRE carriers show presence of RNA foci and poly-GP DPR proteins

Sense and anti-sense RNA foci and DPR proteins have been suggested to represent *C9orf72* HRE-specific gain-of-toxic-function pathological hallmarks [67]. In *post-mortem* CNS of the *C9orf72* HRE carriers, sense and anti-sense RNA foci have been reported in neurons, but also microglia [29, 68]. Notably however, microglia showed RNA foci at a significantly lower frequency than neurons [29]. FISH analysis using a probe against the sense RNA foci demonstrated that RNA foci were present in the nuclei of a subset of the iMG of the *C9orf72* HRE carriers (Fig. 3 a; Supplementary Fig. 9). The antisense foci were not separately assessed.

**Fig. 3.**
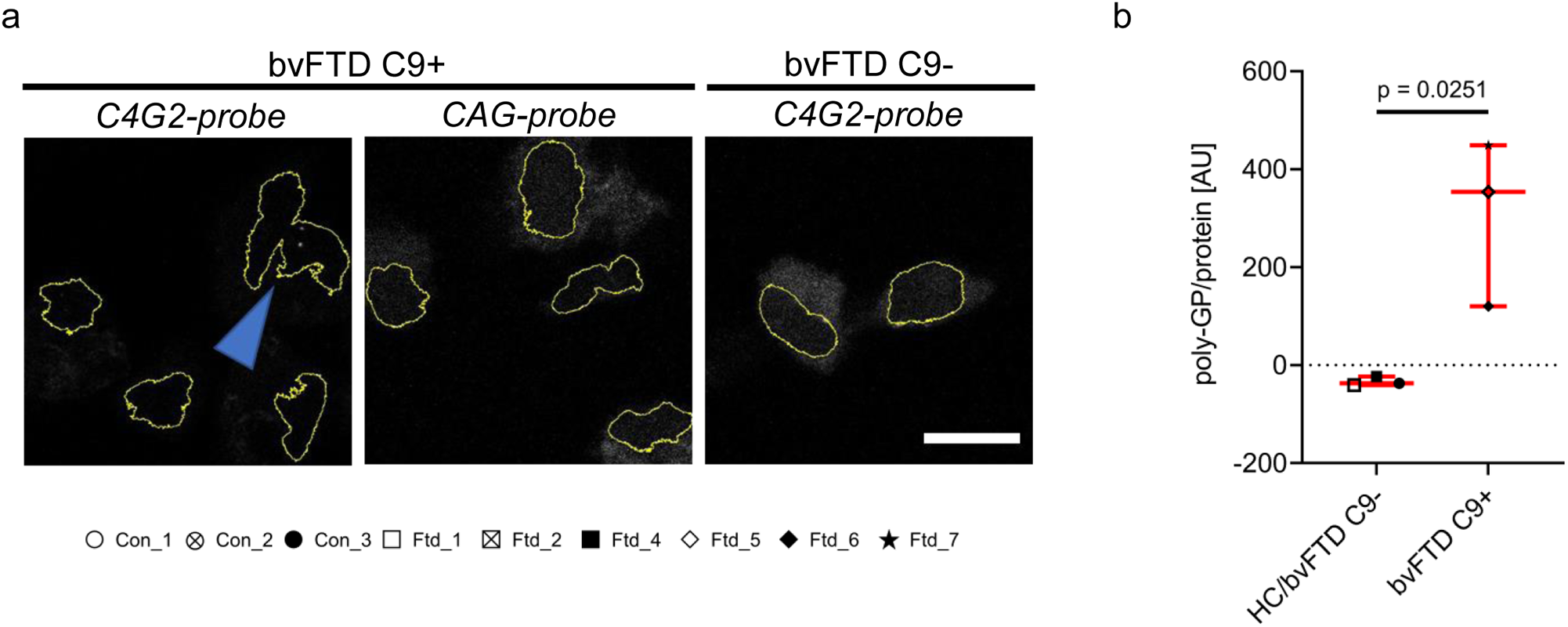
*C9orf72* HRE-derived RNA foci and DPR proteins in iMG from *C9orf72* HRE-carrying bvFTD patients. (a) Intranuclear RNA foci (blue arrow) were detected in iMG of *C9orf72* HRE-carrying bvFTD patients (bvFTD C9+) using fluorescence *in situ* hybridization and a *C9orf72* HRE-targeting probe (*C4G2* probe). No signal was detected in bvFTD C9+ iMG using a probe, which recognizes *CAG* repeats, indicating the specificity of the *C4G2* probe. Also, no signal was detected in the iMG of sporadic bvFTD patients (bvFTD C9-) using the *C4G2* probe. Yellow outlines mark DAPI-positive nuclei. Scale bar = 10 µm. Representative images are shown from two independent differentiation batches. (b) Poly-GP DPR proteins were detected in iMG of C9orf72 HRE-carrying bvFTD patients (bvFTD C9+). Poly-GP levels were measured from one healthy control (HC) iMG line (Con_3) and two sporadic bvFTD patient (bvFTD C9-) iMG lines (Ftd_1, Ftd_4) as *C9orf72* HRE non-carrying controls, and all three bvFTD C9+ iMG lines (Ftd_5, Ftd_6, Ftd_7) from one differentiation batch. Measured poly-GP protein levels were normalized to total protein concentration per sample and were calculated as arbitrary units (AU). Each data point represents the mean value of 3-4 measured samples per line; *p* = 0.0251. The individual iMG lines are indicated by different symbols at the bottom. Descriptive statistics are shown as median ± interquartile range. Two-tailed independent samples t-test was used for statistical analysis. Abbrev.: AU = arbitrary units; bvFTD = behavioral variant frontotemporal dementia; DAPI = 4′,6-diamidino-2-phenylindole; DPR = dipeptide repeat; HC = healthy control; HRE = hexanucleotide repeat expansion; iMG = induced pluripotent stem cell-derived microglia

A sensitive MSD immunoassay against the poly-GP DPR proteins was used to assess whether these were expressed in iMG of *C9orf72* HRE carriers. Poly-GP proteins were specifically detected in the iMG of *C9orf72* HRE carriers but not in the iMG from sporadic bvFTD patient-derived nor control iMG (Fig. 3 b). These findings together indicate that the *C9orf72* HRE-associated gain-of-toxic-function pathological hallmarks, the RNA foci and poly-GP DPR proteins, are present in the iMG of *C9orf72* HRE-carrying bvFTD patients.

### 3.4. iMG of bvFTD patients carrying or not the *C9orf72* HRE do not display TDP-43 pathology

Hyperphosphorylation, C-terminal fragmentation, and cytosolic aggregation of TDP-43 have been generally observed in neurons and glial cells in brain samples of ALS and FTD patients with different genetic backgrounds, including the *C9orf72* HRE carriers [21, 22, 36, 69]. Hence, we investigated whether the iMG of *C9orf72* HRE-carrying and non-carrying bvFTD patients exhibited signs of TDP-43 pathology or expressional changes in TDP-43 levels. According to RNA sequencing, *TARDBP* gene expression was similar between all the three iMG groups (Fig. 4 a). Also, total levels of the full-length TDP-43 proteins were unchanged in all iMG (Fig. 4 b, c). The iMG from either *C9orf72* HRE-carrying or non-carrying bvFTD patients did not show increased phosphorylation of TDP-43 at serine 409/410-1 when compared to control iMG (Fig. 4 b, d). Cleavage of TDP-43 into smaller C-terminal fragments (< 43 kDa) was not observed either in the iMG of *C9orf72* HRE-carrying or non-carrying bvFTD patients (Supplementary Fig. 10). Next, we used our previously published method to quantitatively investigate the translocation of TDP-43 from nucleus to cytosol [51]. iMG from either *C9orf72* HRE-carrying or non-carrying bvFTD patients did not show increased cytosolic-to-nuclear TDP-43 ratio when compared to control iMG (Fig. 4 e, f). Finally, in order to assess at the single cell level whether subpopulations of bvFTD patient iMG showed cytosolic TDP-43 accumulation, the number of iMG with non-pathological *vs.* pathological TDP-43 translocation status (Supplementary Fig. 6 a, b, c) was counted manually. This analysis did not indicate pathological translocation of TDP-43 to the cytosol either. These results collectively indicate that TDP-43 pathology in the form of hyperphosphorylation, C-terminal fragmentation, or cytosolic accumulation is not present in the iMG of bvFTD patients carrying or not the *C9orf72* HRE when compared to the iMG of healthy controls.

**Fig. 4.**
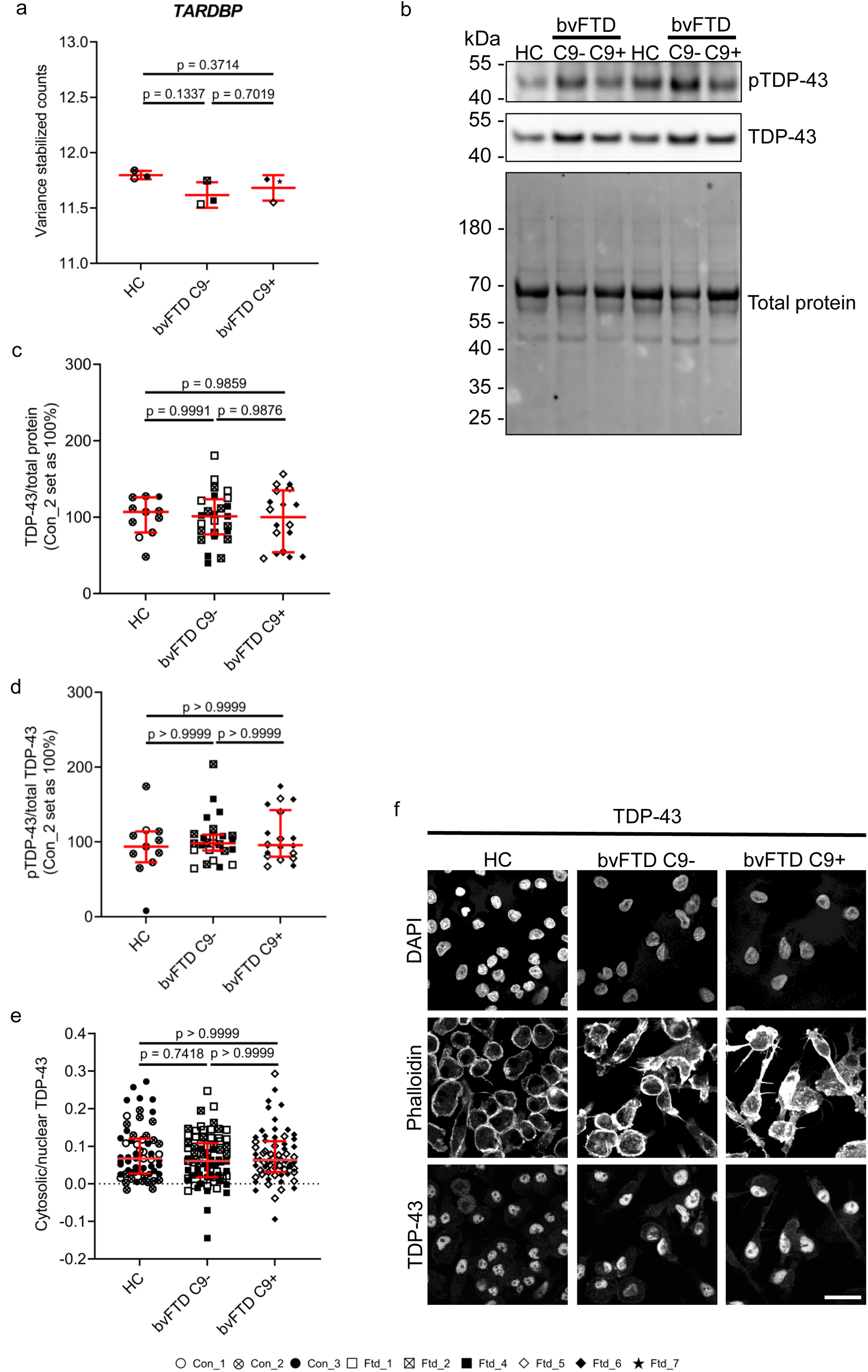
Absence of TDP-43 pathology in iMG from healthy controls, sporadic (C9-) and *C9orf72* HRE-carrying (C9+) bvFTD patients. (a) *TARDBP* gene expression according to RNA sequencing shown for each donor as mean ± standard deviation of variance stabilized counts; *F* = 2.686, *p* = 0.1469; *n* [HC] = 3; *n* [bvFTD C9-] = 3; *n* [bvFTD C9+] = 3; One-way ANOVA followed by Tukey’s multiple comparisons test was used. (b) A representative Western blot image to detect phosphorylated (Ser409/410-1; pTDP-43) and total TDP-43 levels in iMG. (c-d) Quantification of Western blot images regarding total TDP-43 levels normalized to total protein levels (c) and pTDP-43 levels normalized to total TDP-43 levels (d). Each data point represents one biological replicate from three independent differentiation batches in total (c, d). Levels of total TDP-43/total protein (c) or pTDP-43/total TDP-43 (d) were normalized to mean levels of samples from a healthy control (Con_2). For c: *F* = 0.01649, *p* = 0.9837. For d: *H* = 0.3751, *p* = 0.8290. For c, d: *n* [HC] = 11; *n* [bvFTD C9-] = 25; *n* [bvFTD C9+] = 18. (e) Quantification of cytosolic-to-nuclear TDP-43 based on immunofluorescence imaging. Each data point represents one region from three independent differentiation batches. For e: *H* = 1.381, *p* = 0.5012; *n* [HC] = 60; *n* [bvFTD C9-] = 75; *n* [bvFTD C9+] = 66. The individual iMG lines are indicated by different symbols at the bottom. For each group, descriptive statistics are shown as mean ± standard deviation (c) or median ± interquartile range (d-e). One-way ANOVA followed by Tukey’s multiple comparisons test (c) or Kruskal-Wallis test followed by Dunn’s comparisons test (d-e) were used for statistical analysis. (f) Representative microscopy images of TDP-43-stained iMG. DAPI and phalloidin signals were used to outline nuclei and cell boundaries. Scale bar = 30 µm. Abbrev.: bvFTD = behavioral variant frontotemporal dementia; DAPI = 4′,6-diamidino-2-phenylindole; HC = healthy control; HRE = hexanucleotide repeat expansion; iMG = induced pluripotent stem cell-derived microglia; iPSC = induced pluripotent stem cell; *TARDBP* = TAR DNA Binding Protein; TDP-43 = TAR DNA-binding protein 43

### 3.5. iMG of *C9orf72* HRE carriers show changes in proteins involved in autophagosomal-lysosomal pathways

We have previously observed changes in autophagosomal-lysosomal pathways in *C9orf72* HRE-expressing mouse microglial cells [51] and in neuronal cells under *C9orf72* knockdown conditions, mimicking haploinsufficiency [25]. Here, we found that iMG of all three groups showed similar *SQSTM1*, *MAP1LC3B,* and *LAMP2* RNA levels in the RNA sequencing analysis (Fig. 5 a). Protein levels of p62/SQSTM1 and LAMP2-A were also similar in all iMG as indicated by Western blot analysis (Fig. 5 b-d). In addition, no differences in LC3BI to LC3BII conversion, which would indicate alterations in the autophagosomal activity, could be detected between *C9orf72* HRE-carrying or non-carrying bvFTD iMG and control iMG under basal conditions or after Bafilomycin A1 treatment, which induced a similar increase in the LC3BII/LC3BI ratio in all the iMG (Fig. 5 e, f). Quantitative analysis of the immunofluorescence images demonstrated that the number and area of LAMP2-A-positive vesicles as well as LAMP2-A levels within the vesicles were similar between the iMG of sporadic bvFTD and *C9orf72* HRE bvFTD patients. However, these were significantly lower when compared to the iMG of healthy controls (Fig. 6 a-d), suggesting that formation or turnover of the lysosomal vesicles may be altered in the bvFTD iMG from both *C9orf72* HRE carriers and non-carriers.

**Fig. 5.**
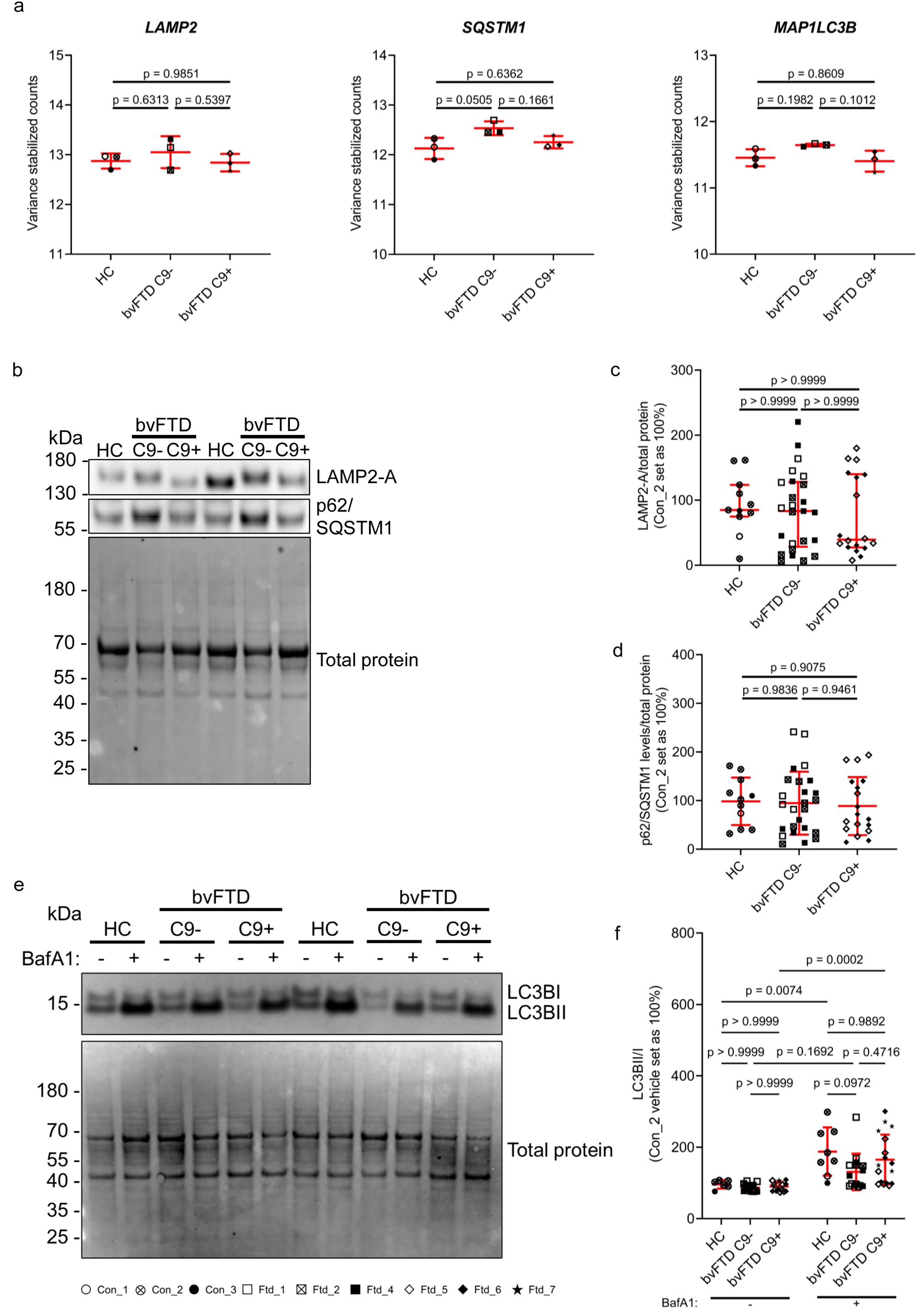
Autophagosomal-lysosomal gene products in iMG from healthy controls, sporadic (C9-) and *C9orf72* HRE-carrying (C9+) bvFTD patients. (a) *LAMP2*, *SQSTM1*, and *MAP1LC3B* gene expression according to RNA sequencing shown for each donor as mean ± standard deviation of variance stabilized counts; *LAMP2*: *F* = 0.7255, *p* = 0.5222; *SQSTM1*: *F* = 4.914, *p* = 0.0545; *MAP1LC3B*: *F* = 3.493, *p* = 0.0986; *n* [HC] = 3; *n* [bvFTD C9-] = 3; *n* [bvFTD C9+] = 3. (b) Representative Western blot images of LAMP2-A, p62/SQSTM1, and total protein signal. Quantification of LAMP2-A (c) and p62/SQSTM1 (d) signals normalized to total protein levels. (e) Representative Western blot images of LC3BI and II with and without Bafilomycin A1 (BafA1) treatment. (f) Quantification of LC3BII/I ratio after normalization to total protein levels. Each data point represents one biological replicate from three (two for e, f) independent differentiation batches in total. For c: *H* = 0.6289, *p* = 0.7302; d: *F* = 0.09748, *p* = 0.9073; f: *F* [treatment] = 37.42, *p* < 0.0001, *F* [group] = 2.655, *p* = 0.0773, *F* [interaction] = 1.459, *p* = 0.2394. For c, d: *n* [HC] = 11; *n* [bvFTD C9-] = 25; *n* [bvFTD C9+] = 18. For f: *n* [HC, vehicle] = 6; *n* [HC, BafA1] = 8; *n* [bvFTD C9-, vehicle] = 14; *n* [bvFTD C9-, BafA1] = 15; *n* [bvFTD C9+, vehicle] = 16; *n* [bvFTD C9+, BafA1] = 17. The individual iMG lines are indicated by different symbols at the bottom. For each group, descriptive statistics are shown as mean ± standard deviation (a, d, f) or median ± interquartile range (c). One-way ANOVA followed by Tukey’s multiple comparisons test (a, d), Kruskal-Wallis test followed by Dunn’s comparisons test (c) or two-way ANOVA followed by Sidak’s multiple comparisons test (f) were used for statistical analysis. Abbrev.: BafA1 = Bafilomycin A1; bvFTD = behavioral variant frontotemporal dementia; HC = healthy control; HRE = hexanucleotide repeat expansion; iMG = induced pluripotent stem cell-derived microglia; kDa = kilo Dalton; LAMP2 = Lysosome-associated membrane glycoprotein 2; *MAP1LC3B*/LC3B = Microtubule-associated proteins 1A/1B light chain 3B; SQSTM1 = Sequestosome 1

**Fig. 6.**
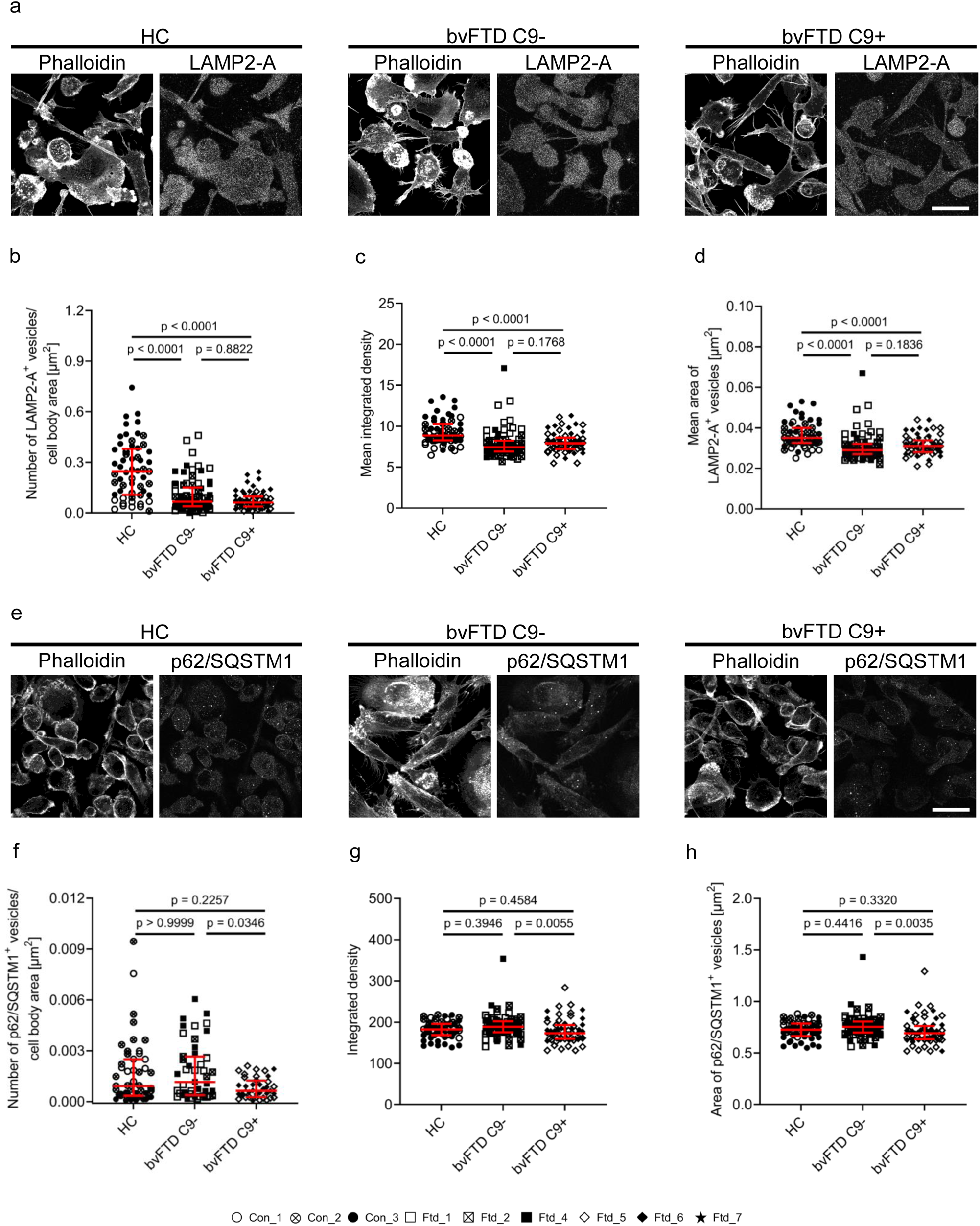
Autophagosomal-lysosomal vesicles in iMG of healthy controls, sporadic (C9-) and *C9orf72* HRE-carrying (C9+) bvFTD patients. Representative immunofluorescence images are shown per group for LAMP2-A (a) and p62/SQSTM1 (e). Phalloidin staining was used to outline the cell boundaries. Scale bars = 30 µm. Each data point represents the mean value (number of vesicles; integrated density and area of LAMP2-A^+^ vesicles) of one region (b, c, d, f) or the value (integrated density and area) of one SQSTM1^+^ vesicle (g, h) from three independent differentiation batches in total. For b: *H* = 48.43, *p* < 0.0001; c: *H* = 39.85, *p* < 0.0001; d: *H* = 40.39, *p* < 0.0001. For b-d: *n* [HC] = 57; *n* [bvFTD C9-] = 78; *n* [bvFTD C9+] = 60. For f: *H* = 6.596, *p* = 0.0370, *n* [HC] = 49; *n* [bvFTD C9-] = 47; *n* [bvFTD C9+] = 39. For g: *H* = 9.761, *p* = 0.0076, *n* [HC] = 53; *n* [bvFTD C9-] = 78; *n* [bvFTD C9+] = 58. For h: *H* = 10.54, *p* = 0.0051, *n* [HC] = 53; *n* [bvFTD C9-] = 78; *n* [bvFTD C9+] = 59. The individual iMG lines are indicated by different symbols at the bottom. For each group, descriptive statistics are shown as median ± interquartile range. Kruskal-Wallis test followed by Dunn’s comparisons test was used for statistical analysis. Abbrev.: bvFTD = behavioral variant frontotemporal dementia; HC = healthy control; HRE = hexanucleotide repeat expansion; iMG = induced pluripotent stem cell-derived microglia; LAMP2 = Lysosome-associated membrane glycoprotein 2; SQSTM1 = Sequestosome 1

TDP-43-negative and p62/SQSTM1-positive cytoplasmic inclusions in the brain have been suggested to represent another hallmark in *C9orf72* HRE carriers [36, 70]. Whereas no p62/SQSTM1-positive inclusions were detected, we observed that iMG from bvFTD patients carrying the *C9orf72* HRE showed fewer p62/SQSTM1-positive vesicles, which were also smaller in size and contained lower p62/SQSTM1 levels, as compared to the iMG from sporadic bvFTD patients but not compared to those from healthy controls (Fig. 6 e-h). These data altogether indicate that iMG of bvFTD patients show a decrease in LAMP2-A-positive lysosomal vesicles, which is further accompanied by reduced p62/SQSTM1-positive vesicles in *C9orf72* HRE-carrying iMG, suggesting potential changes in the autophagosomal-lysosomal pathways.

### 3.6. iMG of *C9orf72* HRE carriers demonstrate alterations in phagocytosis

Several studies have pointed to a potential involvement of C9orf72 in phagocytosis, one of the crucial functions of microglia [39, 66]. Murine C9orf72 localizes in phagolysosomes and late phagosomes in RAW264.7 cells (a murine macrophage cell line) [66]. Moreover, *C9orf72* knockdown has been shown to lead to the upregulation of *Trem2* and *TYRO protein tyrosine kinase-binding protein* in mice [68], which both encode proteins important for the regulation of phagocytosis [71]. Hence, we next characterized the phagocytic activity in the iMG from bvFTD patients carrying or not the *C9orf72* HRE and compared it to that of the control iMG. To this end, we incubated the iMG with pH-sensitive zymosan-coupled bioparticles and followed their uptake. Upon addition of the bioparticles, all iMG first showed increased red fluorescence, indicating phagocytic uptake of the particles, and subsequently increased number and length of motile processes (Fig. 7 a; Supplementary Videos). Area and intensity of the red fluorescence signal were measured over time in live iMG. iMG of bvFTD patients carrying the *C9orf72* HRE showed an increased area (Fig. 7 b, h) and intensity of the fluorescence signal (Fig. 7 c, i) when compared to iMG from healthy controls and sporadic bvFTD patients, suggesting increased phagocytosis of the bioparticles. Serum starvation has been shown to dampen phagocytosis of human iPSC-derived microglia *in vitro* in a previous study [8]. However, in contrast, we observed an increase in the fluorescent area in all iMG after serum starvation compared to the non-starved cells (Fig. 7 d, f, h). Only in the iMG of the sporadic bvFTD patients, the fluorescence intensity was significantly increased after serum starvation compared to the non-starved condition (Fig. 7 e, g, i). Moreover, iMG of sporadic bvFTD patients also showed increased area (Fig. 7 d, f, h) and intensity (Fig. 7 e, g, i) of the fluorescent signal as compared to healthy controls under serum starvation. The iMG of bvFTD patients carrying the *C9orf72* HRE still showed the biggest increase in the fluorescent area and intensity as compared to the other iMG also after serum starvation, further supporting the notion that they show increased phagocytic activity. These results suggest that iMG carrying the *C9orf72* HRE display increased phagocytosis both under basal and serum-starved conditions compared to the iMG from sporadic bvFTD patients and healthy controls. After serum starvation, also the iMG of sporadic bvFTD patients show increased phagocytosis compared to the iMG from the healthy controls.

**Fig. 7.**
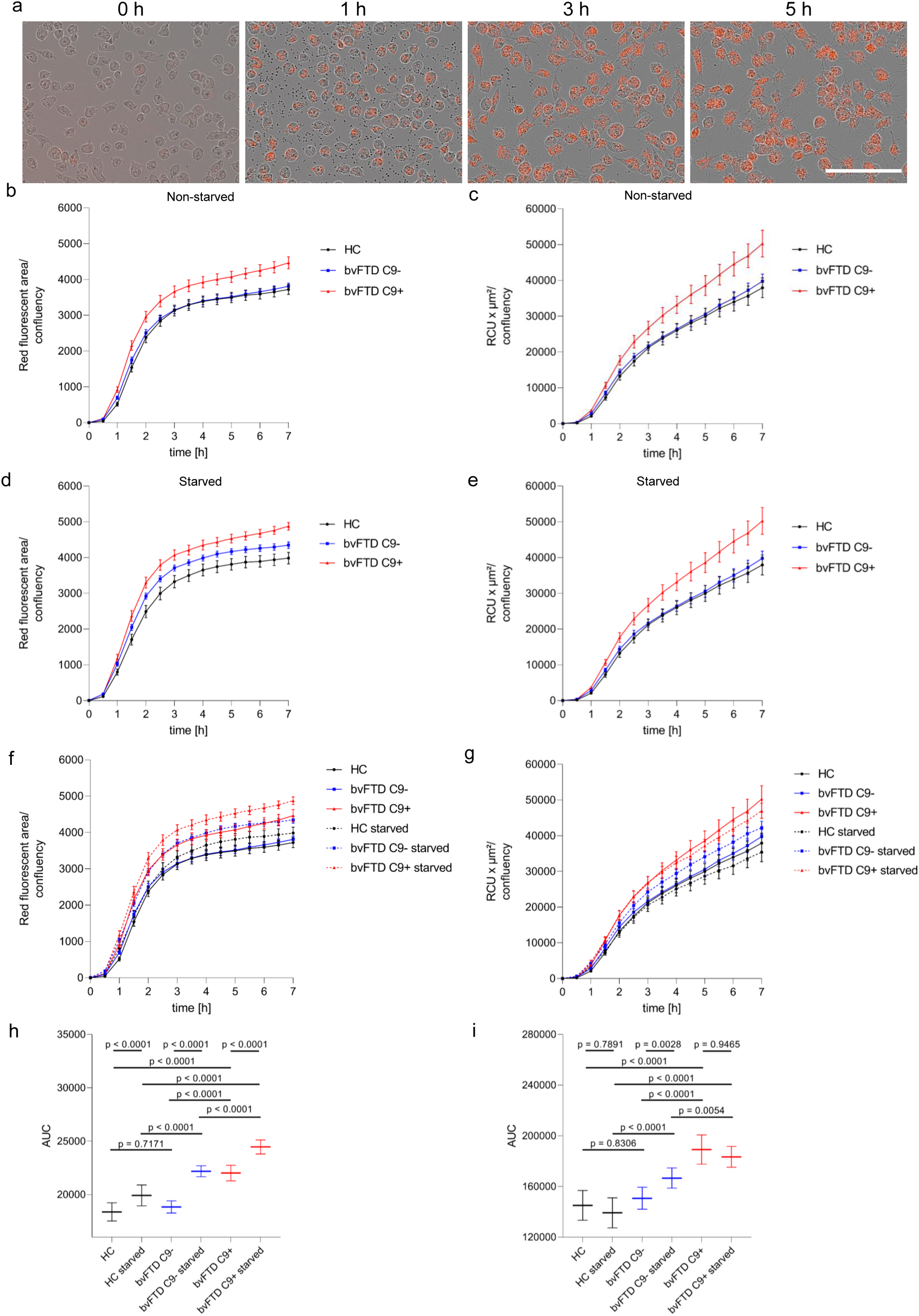
Phagocytosis in iMG of healthy controls, sporadic (C9-) and *C9orf72* HRE-carrying (C9+) bvFTD patients. iMG cultured under normal conditions were incubated with pHrodo zymosan beads and imaged over time. (a) Representative images of iMG from a healthy donor. Scale bar = 150 µm. For each well, red fluorescence was measured over time and normalized to confluency prior to zymosan bead addition. Area of red fluorescence (b, d, f, h) or fluorescence intensity (c, e, g, i) per well were calculated. Data are shown as mean ± standard error of mean (b-g) or mean ± standard deviation (h, i). Statistical testing (one-way ANOVA followed by Sidak’s multiple comparisons test) was applied after area under the curve (AUC) analysis (h: *F* = 97.11, *p* < 0.0001, i: *F* = 38.34, *p* < 0.0001). Data are shown from two independent differentiation batches (*n* [HC] = 12; *n* [bvFTD C9-] = 12; *n* [bvFTD C9+] = 7; *n* [bvFTD C9+ starved] = 8). Abbrev.: bvFTD = behavioral variant frontotemporal dementia; HC = healthy control; HRE = hexanucleotide repeat expansion; iMG = induced pluripotent stem cell-derived microglia; RCU = red calibrated unit

### 3.7. iMG of *C9orf72* HRE carriers exhibit expressional changes in genes related to phagocytosis and autophagy

Next, we investigated whether genes related to phagocytosis and autophagy were differentially expressed in the iMG of *C9orf72* HRE bvFTD patients. According to RNA sequencing, no differences in the expression of receptors, which have been shown to bind zymosan, including *CLEC7A* or *TLR2*, *5*, and *6* [72, 73], could be detected (Supplementary Fig. 11 a). Pathway analysis of genes involved in phagocytosis regulation and recognition and autophagy showed that *CD93*, Ras-related C3 botulinum toxin substrate 2 (*RAC2*), *CD300A,* and Coronin 1C (*CORO1C*) were expressed at higher levels in the iMG carrying the *C9orf72* HRE as compared to iMG from healthy controls or sporadic bvFTD patients (Supplementary Fig. 11 b). However, these changes were significant only in comparison to sporadic bvFTD iMG but not to the iMG from healthy controls (Supplementary Table 2). Similarly, the expression of heat shock protein 90 alpha family class A member 1 (*HSP90AA1*) and tubulin alpha 1b (*TUBA1B*), which are related to autophagy, were significantly upregulated in the iMG of *C9orf72* HRE carriers as compared to iMG from sporadic bvFTD patients but not to iMG from healthy controls (Supplementary Fig. 11 a; Supplementary Table 2). Thus, these RNA-based analyses of genes regulating autophagy or phagocytosis do not indicate similar changes in the iMG of healthy controls and sporadic bvFTD patients, which compared to the expressional changes in the iMG of *C9orf72* HRE-carrying bvFTD patients would explain the observed increased phagocytic activity in the iMG of *C9orf72* HRE-carrying bvFTD patients.

### 3.8. iMG of *C9orf72* HRE carriers and sporadic bvFTD patients show differential expression of genes involved in processes of RNA and protein regulation and cellular energy metabolism

Gene-set enrichment analysis revealed that pathways involved in the regulation of different RNA (tRNA, rRNA, mRNA, ncRNA) processes and protein translation, localization, and degradation were specifically upregulated in the iMG from the *C9orf72* HRE-carrying bvFTD patients as compared to sporadic bvFTD patients. Furthermore, genes involved in energy metabolism, including mitochondrial metabolism, were upregulated in the iMG from *C9orf72* HRE carriers as compared to sporadic bvFTD patients. Interestingly, the iMG of sporadic bvFTD patients showed a decreased expression pattern in pathways related to rRNA, tRNA, and ncRNA metabolism when compared to iMG of healthy controls (Fig. 8).

**Fig. 8.**
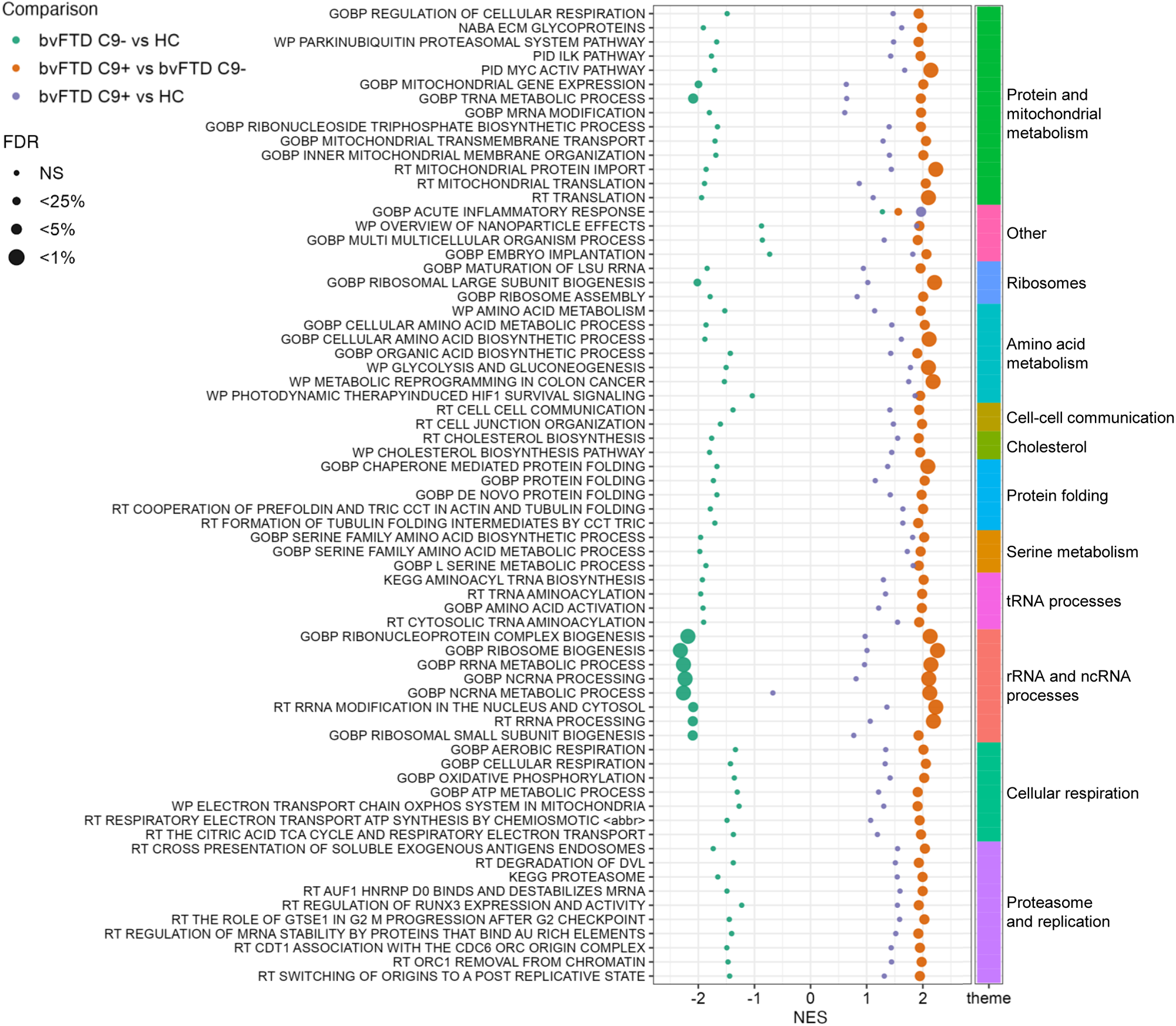
Differentially enriched pathways in iMG of healthy controls, sporadic (C9-), and *C9orf72* HRE-carrying (C9+) bvFTD patients. The dot plot displays, for the three differently colored comparisons, the gene set enrichment analysis (GSEA) results for the MSigDB gene set collections GO Biological Processes (C5 GO BP) and Canonical Pathways (C2 CP). Normalized enrichment score (NES) is shown in the x-axis whereas the categorical statistical significance (FDR) is indicated by the size of the marker. Data are shown from one differentiation batch. Abbrev.: bvFTD = behavioral variant frontotemporal dementia; *CLEC7A* = C-type lectin domain containing 7A; HC = healthy control; HRE = hexanucleotide repeat expansion; iMG = induced pluripotent stem cell derived microglia; NES = Normalized Enrichment Score; NS = not significant

### 3.9. iMG of *C9orf72* HRE carriers display differential expression of genes involved in acute inflammatory response

In comparison to healthy controls, iMG from *C9orf72* HRE bvFTD patients specifically showed a higher expression of genes involved in acute inflammatory response, namely orosomucoid 1 (*ORM1*), prostaglandin-endoperoxide synthase 2 (*PTGS2*), and S100 calcium-binding protein A8 (*S100A8*) (Fig. 8; Supplementary Figure 12 a, Supplementary Table 2). The levels of some cytokines and chemoattractants related to microglial activation, such as C-C motif chemokine ligand 2 (CCL2), chitinase 1 (CHIT1), IL-18, and tumor necrosis factor α (TNFα), have been found elevated in the plasma or cerebrospinal fluid (CSF) of *C9orf72* HRE patients, whereas the levels of C-X-C motif chemokine ligand 10 (CXCL10) were decreased. The levels of IL-1β also negatively correlated with survival [74–77]. However, the RNA levels of none of these genes were altered in the *C9orf72* HRE-carrying or sporadic bvFTD patient iMG as compared to control iMG. An exception was *IL1B*, which showed higher levels in the iMG of *C9orf72* HRE bvFTD patients as compared to sporadic bvFTD patients but not as compared to healthy controls (Supplementary Fig. 12 b; Supplementary Table 2). Finally, heterozygous and homozygous *3110043O21Rik* (mouse *C9orf72*) knockout mice were recently reported to exhibit an increase in interferon response microglia (IRM) signature and a decrease in gene expression linked to active response microglia (ARM). In line with these changes observed in mice, pronounced IFN type I signaling in the peripheral mononuclear blood cells, monocyte-derived macrophages, and cerebellar tissue of *C9orf72* HRE-carrying ALS patients when compared to sporadic ALS patients were reported [39, 78]. However, in the present study, no obvious changes in the IRM or ARM signature could be detected in the iMG of *C9orf72* HRE-carrying bvFTD patients when compared to the iMG from sporadic bvFTD patients or healthy controls according to our RNA sequencing data (Supplementary Table 2).

## 4. Discussion

In this study, we have generated for the first time iMG from *C9orf72* HRE-carrying and sporadic bvFTD patients and compared them to iMG from healthy controls. Using RNA sequencing, we confirmed that all the iMG similarly expressed typical microglial genes, indicating that they successfully acquired microglial identity. This was further confirmed by immunofluorescence staining of several microglial markers at the protein level. These findings validate the applicability of a previously published protocol [13] to differentiate iPSCs into iMG *in vitro* from both *C9orf72* HRE-carrying and sporadic patients clinically diagnosed with bvFTD.

Our examinations indicated that *C9orf72* HRE-carrying iMG from bvFTD patients did not present reduced C9orf72 levels but showed formation of sense RNA foci in the nuclei and expression of poly-GP DPR proteins, suggesting that under basal culture conditions, the iMG partially recapitulate the *C9orf72* HRE-derived pathological hallmarks typically detected in the CNS of the patients. These findings are in accordance with previous studies on iMG from *C9orf72* HRE ALS and ALS/FTD patients, showing the presence of RNA foci and DPR proteins [41, 42]. Interestingly, in the study by Vahsen et al., poly-GA and poly-GP proteins were detected at higher levels in iPSC-derived motoneurons compared to iMG [42]. Reduced C9orf72 protein levels were detected in two studies when comparing the *C9orf72* HRE iMG to healthy control [41, 42] or to isogenic control lines in one study (Banerjee et al., 2023). Previously, we have detected RNA foci, but not DPR proteins nor *C9orf72* haploinsufficiency, in *C9orf72* HRE-carrying skin biopsy-derived fibroblasts [43] from which the iPSCs and iMG used in this study were generated. These findings may implicate that specific mechanisms, such as methylation of the *C9orf72* promoter [79] involved in the regulation of the *C9orf72* expression, may differ from those in the other cell types or the brain where haploinsufficiency has been detected.

The iMG from sporadic or *C9orf72* HRE-carrying bvFTD patients did not display signs of TDP-43 pathology as indicated by the absence of clear cytoplasmic TDP-43 inclusions, TDP-43 hyperphosphorylation, cleavage to C-terminal fragments, or increased translocation of TDP-43 from nucleus to cytosol. No TDP-43 pathology in the fibroblasts of the bvFTD patients carrying or not the *C9orf72* HRE compared to the fibroblasts from healthy donors was detected in our previously published study either [43]. In our previous studies using mouse microglial BV-2 cells overexpressing the *C9orf72* HRE, we found that the cells expressed poly-GA and poly-GP DPR proteins but there were no RNA foci nor changes in the C9orf72 levels. These cells showed slightly increased nucleus-to-cytoplasm translocation of TDP-43, suggesting mild TDP-43 pathology [51]. Altogether, these findings are in agreement with studies by others and support the idea that cytosolic TDP-43 accumulation and formation of TDP-43 pathology might be driven by accumulation of DPR proteins [80] rather than the presence of RNA foci or *C9orf72* haploinsufficiency. Similarly to the present study, the iMG from *C9orf72* HRE ALS and ALS/FTD patients expressing poly-GA and poly-GP proteins also lacked TDP-43 pathology in previous studies [41, 42]. Therefore, these studies could indicate that development of TDP-43 pathology in the iMG might need a longer time or require the presence of additional stress. Early nuclear TDP-43 pathology detectable with TDP-43 RNA aptamers but not with classic antibodies has been suggested to occur prior to cytoplasmic accumulation of TDP-43 and precede clinical symptoms in ALS patients [81]. Testing whether similar nuclear TDP-43 pathology is present in the bvFTD patient-derived iMG might give further insights into whether the iMG recapitulate an early stage of TDP-43 pathology.

We have reported that p62/SQSTM1-positive vesicles were enlarged and accumulated in the fibroblasts of bvFTD patients compared to healthy controls regardless of the presence or absence of the *C9orf72* HRE [43]. p62/SQSTM1 protein localizes in the autophagosomes and undergoes autophagosomal degradation. Therefore, alterations in its levels may indicate changes in the autophagosomal activity [82]. No changes in the autophagosomal function in the bvFTD patient fibroblasts were detected in our previous study [43]. Here, we detected no differences in LC3BI to LC3BII conversion in *C9orf72* HRE-carrying or sporadic bvFTD iMG and control iMG under basal conditions or following Bafilomycin A1 treatment, indicating unaltered autophagosomal activity. However, in contrast to the fibroblasts, immunofluorescence studies revealed a reduced number and size of p62/SQSTM1-positive vesicles in the iMG of *C9orf72* HRE-carrying bvFTD patients, which might be indicative of enhanced function of the autophagosomal pathway. Furthermore, regardless of the *C9orf72* HRE carriership, iMG of all bvFTD patients displayed a lower number of LAMP2-A-positive vesicles, which also showed decreased size and levels of LAMP2-A, suggesting possible alterations in lysosomes. Reduced number of LAMP2-positive vesicles in iPSC-derived motoneurons from sporadic ALS and *C9orf72* HRE-carrying ALS patients compared to healthy control has also previously been reported [30, 83]. Together, these findings suggest that there may be cell type-specific alterations in the autophagosomal-lysosomal pathways in the bvFTD patient-derived cells, including microglia, but detailed characterization of such potential changes remains the subject of further studies.

Autophagosomal-lysosomal function is intimately linked to phagocytosis in phagocytosing cells, such as microglia. Increased phagocytosis and synaptic pruning have been shown to take place in microglia of homozygous *C9orf72* knock-out mice [39]. Here, iMG of *C9orf72* HRE-carrying bvFTD patients showed significantly increased fluorescence signal of phagocytosed zymosan beads, implicating their increased uptake by phagocytosis. In a similar manner, Lorenzini et al. reported increased Aβ but not synaptoneurosome uptake [41], suggesting that the *C9orf72* HRE-carrying microglia may show differences in the phagocytosis of different substances. In the study by Banerjee et al., on the other hand, reduced phagocytosis of zymosan beads was detected with 10-fold higher concentrations of the beads as compared to our study and this reduction was observed already after 30 to 75 min [23]. This difference might be explained by the prominently lower C9orf72 levels [23] as compared to the *C9orf72* HRE iMG in our study. They also detected disrupted autophagy initiation after Bafilomycin A1 treatment, but no differences in the number of p62/SQSTM1 vesicles under basal conditions as compared to isogenic controls [23]. Notably, in our study, there was a decrease in the number, intensity, and size of the in p62/SQSTM1 vesicles between the iMG from *C9orf72* HRE *vs.* sporadic bvFTD patients.

These functional findings together with lower numbers of p62/SQSTM1- and LAMP2-A-positive vesicles, possibly indicating their increased turnover, in *C9orf72* HRE-carrying iMG suggest that these cells intrinsically show an increased phagocytic activity. Interestingly, serum starvation led to a higher phagocytic activity in *C9orf72* HRE-carrying iMG but also in the iMG of sporadic bvFTD patients. Thus, a fewer number of LAMP2-A-positive vesicles and a higher phagocytic activity under serum starvation as compared to iMG of the healthy controls suggest potential changes in the autophagosomal-lysosomal pathway also in sporadic bvFTD iMG under specific conditions. According to our RNA sequencing data, no specific genes or pathways related to phagocytosis and autophagy showing differential expression in both iMG from healthy controls and sporadic bvFTD patients compared to the iMG from *C9orf72* HRE-carrying bvFTD patients could be identified, which could explain the observed augmented phagocytic activity of the *C9orf72* HRE-carrying iMG. Therefore, a proteomic approach might be useful to further investigate the underlying mechanisms of the increased phagocytosis and possible alterations in the autophagosomal and lysosomal pathways. The severity of impaired phagocytosis of iMG from ALS patients has been correlated with disease progression pointing to a potential clinical relevance of this important microglial function [84].

According to gene-set enrichment analysis, the RNA profiles of iMG of bvFTD patients carrying the *C9orf72* HRE *vs.* sporadic bvFTD patients showed remarkable differences. These differentially enriched genes and pathways point towards strong differences in processes related to RNA and protein regulation and energy metabolism. In general, these pathways showed an upregulation in the *C9orf72* HRE-carrying iMG as compared to the sporadic bvFTD iMG. However, in accordance with the findings by Lorenzini et al., the RNA profiles of the *C9orf72* HRE-carrying iMG modestly differed from those of the healthy control iMG [41]. In contrast, the sporadic iMG showed marked differences to the healthy control iMG in their RNA profiles, indicated by downregulation of *e.g.*, pathways related to RNA and protein regulations, suggesting an opposite profile to that in the *C9orf72* HRE patient iMG. In the future, more detailed investigations will be needed to investigate which kind of functional consequences these differences in the gene profiles may have and if they are further reflected in other cell types of the same patients as well. Elucidating whether functional changes in these essential cellular processes in the bvFTD patient-derived iMG are indeed present might prove important *e.g.,* in future drug design or biomarker discovery.

Even though several inflammation-related proteins have been reported to be differentially expressed in the CSF or plasma of *C9orf72* HRE-carrying patients, we did not observe an overt inflammatory phenotype in the iMG from bvFTD patients carrying or not the *C9orf72* HRE under basal conditions. Neither did the RNA sequencing reveal a phenotypic change of the bvFTD iMG to the recently reported ARM or IRM [39, 78]. However, compared to the iMG from healthy controls, iMG from *C9orf72* HRE-carrying bvFTD patients did show an increased expression of *ORM1*, *PTGS2*, and *S100A8*, which are genes involved in acute inflammatory response. *ORM1* gene expression was also increased in the iMG of sporadic bvFTD patients as compared to healthy control iMG, but the expression of *PTGS2* and *S100A8* was only elevated in *C9orf72* HRE carriers compared to the other two groups. The expression of *PTGS2*, also known as cyclooxygenase-2 (*COX-2*), can be regulated by S100A8, and S100A8 has been shown to induce the release of pro-inflammatory cytokines by mouse microglia *in vitro* [85, 86]. Therefore, stimulation the *C9orf72* HRE iMG with different inflammatory stimuli and defining their RNA profiles or measuring the release of inflammatory cytokines might provide further insights into whether these cells show differential activation states or function as compared to the iMG from sporadic bvFTD patients or healthy controls. Furthermore, since none of the proteins encoded by the previously mentioned genes were significantly differentially expressed in the brains of sporadic *vs. C9orf72* HRE-carrying ALS patients in a previous report [87], it might be interesting to measure their levels in *e.g.* blood samples or in CSF of bvFTD patients to evaluate if these RNA-level findings could be translated to potential biomarkers.

One limitation of our study is the use of monocultures of the iMG. Previous studies have indicated that microglia readily respond to the signals coming from their environment and surrounding cells, such as neurons, and that their activation stage changes upon these cues. Moreover, isolated microglia or microglia cultured *in vitro* show different RNA expression profiles than the brain-resident microglia *in vivo* [88]. In line with these studies, several mouse models of the *C9orf72* HRE suggest that neuron-microglia interaction might cause microglia activation and affect their function [89]. Thus, it would be important to study also the *C9orf72* HRE-carrying iMG, for example, in co-cultures with neurons. This might be especially important since human iPSC-derived microglia-neuron co-cultures show higher expression of *C9orf72* as compared to monocultures of microglia [14]. A higher *C9orf72* gene expression might also lead to a stronger presence of *C9orf72* HRE-derived pathological hallmarks and therefore more pronounced disease phenotypes in microglia, and potentially subsequently in neurons as well. Thus, a neuron-microglia co-culture approach might give deeper insights into disease pathogenesis and underlying mechanisms. Since co-culturing the *C9orf72* HRE-carrying ALS iMG with motoneurons was shown to increase neuronal vulnerability [23, 42], it would be important to assess in the future whether this also applies to co-cultures with neurons from bvFTD patients. On the other hand, we observed that the iMG of bvFTD patients in monocultures showed differences in potential disease-relevant pathways, such as phagocytosis and RNA, protein, and energy metabolism, as compared to controls. Moreover, the iMG of *C9orf72* HRE carriers showed formation of RNA foci and poly-GP DPR proteins, partially recapitulating the pathological changes in the patient brain. These findings suggest that despite representing a likely immature and simplified model system, monocultures of iMG may provide important initial insights into potential microglial alterations in bvFTD that may then be further examined in other more complex and sophisticated disease models and even in human brain. In the future, it would add to the translational value of the iMG if the alterations found in this study could be confirmed in the microglia in the brains of the *C9orf72* HRE-carrying patients. Also, isogenic control lines of *C9orf72* HRE-carrying iPSCs would be useful as additional controls in addition to the healthy control lines. This might enable identification of the pathways that are altered specifically due to the *C9orf72* HRE. Moreover, targeting the formation of RNA foci and DPR proteins through antisense oligonucleotides might shed light onto the exact underlying molecular mechanisms of the observed phenotypes of the *C9orf72* HRE-carrying iMG. Finally, since high variance was detected in some experiments, inclusion of a larger number of iMG lines might increase the statistical power in future experiments.

Taken together, here we report for the first time RNA profiles and cell pathological and functional phenotypes of the iMG derived from both *C9orf72* HRE-carrying and non-carrying sporadic bvFTD patients. iMG from the *C9orf72* HRE carriers displayed the presence of RNA foci and DPR proteins and decreased number and size of p62/SQSTM1- and LAMP2-A-positive vesicles, which coincided with increased phagocytosis, suggesting alterations in the autophagosomal-lysosomal pathways. The *C9orf72* HRE-carrying bvFTD iMG also specifically showed differential expression of genes involved in RNA, protein, and energy metabolism. Furthermore, increased phagocytosis in sporadic bvFTD iMG after serum starvation as well as similarly decreased number and size of the LAMP2-A-positive vesicles to the *C9orf72* HRE-carrying iMG shed light for the first time on partially common alterations of bvFTD iMG regardless of the genetic background. The present study represents an initial starting point for using patient-derived iMG as a disease model for bvFTD. Future investigations utilizing different iMG-based model systems will help to better understand the underlying molecular mechanisms of different clinical and genetic forms of FTD and the involvement of microglia in the disease pathogenesis.

## Abbreviations

ab: Antibody
*AIF1*: Allograft inflammatory factor 1
ALS: Amyotrophic lateral sclerosis
ANOVA: Analysis of variance
ARM: Active response microglia
AUC: Area under the curve
BSA: Bovine serum albumin
bvFTD: Behavioral variant frontotemporal dementia
*C1QA*: Complement C1q subcomponent subunit A
*C9orf72*: Chromosome 9 open reading frame 72
*CCL2*: C-C motif chemokine ligand 2
CD: Cluster of differentiation
*CHIT1*: Chitinase 1
CNS: Central nervous system
*CORO1C*: Coronin 1C
*COX-2*: Cyclooxygenase-2
CSF: Cerebrospinal fluid
CX3CR1: CX3C chemokine receptor 1
*CXCL10*: C-X-C motif chemokine ligand 10
DAPI: 4′,6-diamidino-2-phenylindole
DMSO: Dimethyl sulfoxide
DNA: Deoxyribonucleic acid
DPBS: Dulbecco’s phosphate-buffered saline
DPR: Dipeptide repeat
EDTA: Ethylenediaminetetraacetic acid
FBS: Fetal bovine serum
FISH: Fluorescence *in situ* hybridization
FTD: Frontotemporal dementia
*GAS6*: Growth arrest-specific 6
*GPR34*: G-protein coupled receptor 34
HC: Healthy control
*HEXB*: Beta-hexosaminidase subunit beta
HRE: Hexanucleotide repeat expansion
*HSP90AA1*: Heat shock protein 90 alpha family class A member 1
IBA1: Ionized calcium binding adapter molecule 1
IL: Interleukin
iMG: iPSC-derived microglia
iPSC: induced pluripotent stem cell
IRM: Interferon response microglia
kDa: kilo Dalton
LAMP: Lysosome-associated membrane glycoprotein
*MAP1LC3B*/LC3B: Microtubule-associated proteins 1A/1B light chain 3B
MDMi: monocyte derived microglia
*MERTK*: MER Proto-Oncogene, Tyrosine Kinase
MSD: Meso Scale Discovery
*NANOG*: Nanog homeobox
*ORM1*: Orosomucoid 1
P2RY12: P2Y purinoceptor 12
PCR: Polymerase chain reaction
PDL: Poly-D-lysine
*POU5F1*: POU domain, class 5, transcription factor 1
*PROS1*: Protein S
*PSEN2*: Presenilin 2
*PTGS2*: Prostaglandin-endoperoxide synthase 2
*RAC2*: Ras-related C3 botulinum toxin substrate 2
RCU: Red calibrated unit
RNA: Ribonucleic acid
RT: Room temperature
*S100A8*: S100 calcium-binding protein A8
SDS: Sodium dodecyl sulphate
*SOD1*: Superoxide dismutase 1
*SOX2*: SRY (sex determining region Y)-box 2
SQSTM1: Sequestosome 1
SSEA4: Stage-specific embryonic antigen 4
*TARDBP*: TAR DNA Binding Protein
TDP-43: TAR DNA-binding protein 43
TMEM119: Transmembrane protein 119
*TNF*: Tumor necrosis factor
TRA181: Tumor-related Antigen-1-81
TREM2: Triggering receptor expressed on myeloid cells 2
*TUBA1*: Tubulin Alpha 1b
*TYROBP*: TYRO protein tyrosine kinase-binding protein
UMAP: Uniform Manifold Approximation and Projection
WB: Western blotting

## Declarations

### Consent for publication

Not applicable

### Authors’ contributions

Annakaisa Haapasalo and Hannah Rostalski conceptualized the design of the study. Anne M. Portaankorva, Eino Solje, Kasper Katisko, Päivi Hartikainen, Šárka Lehtonen, Henna Martiskainen, and Jari Koistinaho collected and processed patient samples and data. Nadine Huber, Stina Leskelä, Hannah Rostalski, Petra Mäkinen, Sohvi Ohtonen, and Henna Jäntti validated the methods. Šárka Lehtonen, Teemu Natunen, Henna Martiskainen, Mari Takalo, Tarja Malm, Annakaisa Haapasalo, and Mikko Hiltunen supervised the study. Hannah Rostalski, Dorit Hoffmann, and Tomi Hietanen performed cell culture experiments. Hannah Rostalski, Tomi Hietanen, Dorit Hoffmann, Nadine Huber, and Šárka Lehtonen performed biochemical analyses and RNA extraction. Hannah Rostalski performed live cell analysis. Ashutosh Dhingra and Salvador Rodriguez-Nieto performed MSD immunoassays. Hannah Rostalski, Tomi Hietanen, and Viivi Pekkala analyzed microscopy images. Mari Takalo, Henna Martiskainen, and Mikko Hiltunen provided previous RNA sequencing data. Teemu Kuulasmaa and Sami Heikkinen performed analysis on RNA sequencing data. Hannah Rostalski performed all other analyses and data presentation. Hannah Rostalski wrote the initial draft of the article and together with Tomi Hietanen revised the manuscript. Hannah Rostalski, Tomi Hietanen, Sami Heikkinen, and Šárka Lehtonen prepared the figures. All authors have contributed to revising the manuscript draft. All authors have read and approved the final manuscript.

## Acknowledgements

We would like to thank all the study participants for their valuable blood and skin biopsy donations. Thank you to Laila Kaskela and Eila Korhonen for iPSCs generation and Jenni Voutilainen and Ida Hyötyläinen for assistance characterizing the generated iPSC lines. H.R., S.L., N.H., and S.O. are or have been supported by the University of Eastern Finland (UEF) Doctoral Programme in Molecular Medicine (DPMM). H.R. has been supported by the GenomMed doctoral programme. UEF Cell and Tissue Imaging Unit, supported by Biocenter Kuopio and Biocenter Finland is acknowledged for providing IncuCyte® S3 and LSM800 training and facilities. UEF Bioinformatics Center is acknowledged for the use of the high-performance computing cluster. This publication is part of a project that has received funding from the European Union’s Horizon 2020 research and innovation programme under the Marie Skłodowska-Curie grant agreement no 740264. This study is part of the research activities of the Finnish FTD Research Network (FinFTD).

This study was supported by academic research grants from the Academy of Finland, grant nos. 315459 (A.H.), 315460 (A.M.P.), 288659 (T.N.), 328287 (T.M.), 330178 (M.T.) and 338182 (M.H.); Yrjö Jahnsson Foundation, grant no. 20187070 (A.H.); Päivikki and Sakari Sohlberg Foundation (A.H.); ALS tutkimuksen tuki ry. registered association (H.R., S.L., N.H.); The Maud Kuistila Memorial Foundation (H.R.); Orion Research Foundation sr (H.R., E.S.); The Finnish Cultural Foundation (T.H.); Kuopio University Foundation (H.R.); Sigrid Jusélius Foundation (A.H., M.H., E.S., Š.L.); The Finnish Brain Foundation (E.S., K.K.); Instrumentarium Science Foundation (E.S.); The Finnish Medical Foundation (E.S., K.K.); Maire Taponen Foundation (K.K.); The Strategic Neuroscience Funding of the University of Eastern Finland (A.H., M.H.); and Neurocenter Finland – AlzTrans pilot project (M.H.).

## Data Availability

The datasets used and/or analyzed during the current study are available from the corresponding author on reasonable request.

## Ethics approval

The patient data used in this project is pseudonymized and is handled using code numbers only. Handling of cell cultures related to microglia differentiation is performed with the permission 123/2016 from the Research Ethics Committee of the Northern Savo Hospital District. The study was performed in accordance with the principles of the declaration of Helsinki.

## Consent to participate

Donors of skin biopsies have given their written informed consent prior to sample collection.

## Supplementary Information (SI)

**Supplementary Table 1.**
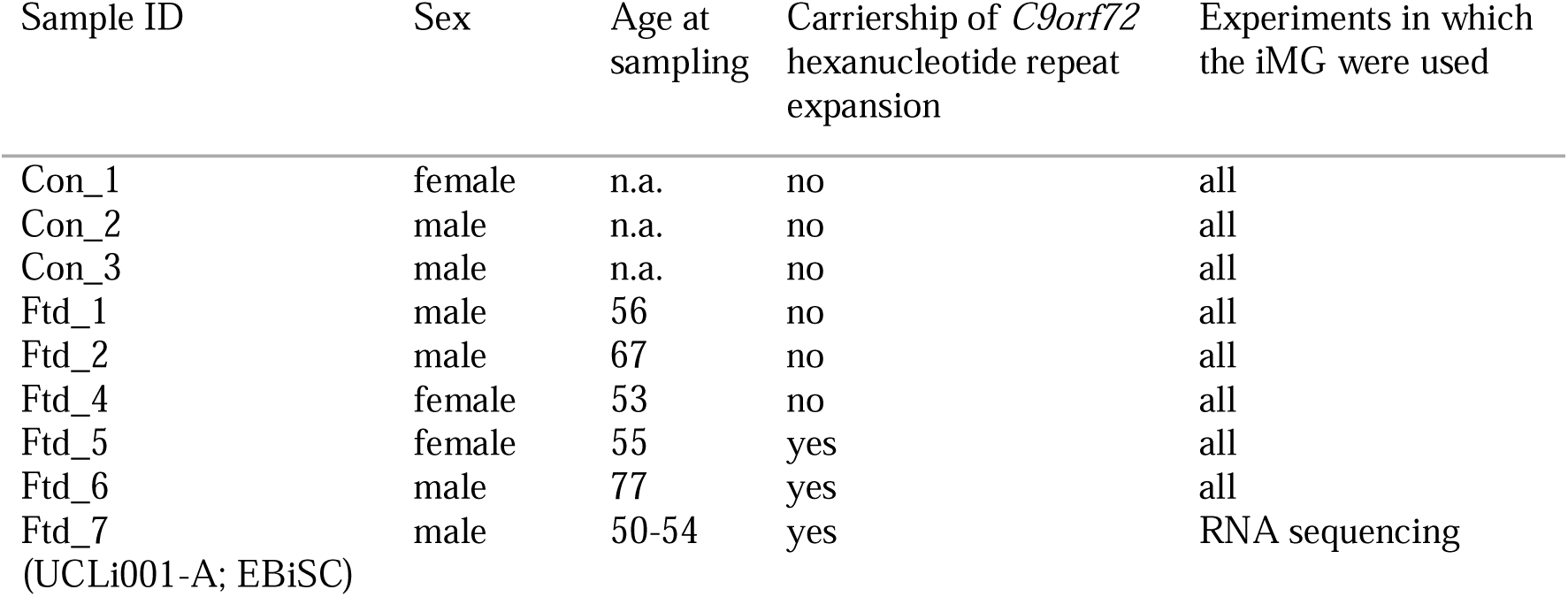
Information on donors; n.a. = not available.

**Supplementary Table 2. DEG and GSEA results on RNA sequencing data (s. online)**

**Supplementary Fig. 1. iPSCs from healthy controls, sporadic (C9-) and *C9orf72* HRE-carrying (C9+) bvFTD patients express pluripotency genes.** qPCR was used to assess gene expression levels of different pluripotency marker genes in iPSCs. Each datapoint indicates technical replicates in each iPSC line. Abbrev.: bvFTD = behavioral variant frontotemporal dementia; HC = healthy control; HRE = hexanucleotide repeat expansion; iPSCs = induced pluripotent stem cells; *NANOG* = Nanog homeobox; *POU5F1* = POU domain, class 5, transcription factor 1; *SOX2* = SRY (sex determining region Y)-box 2

**Supplementary Fig. 2. iPSCs from healthy controls, sporadic (C9-) and *C9orf72* HRE-carrying bvFTD (C9+) patients express pluripotency markers**. iPSCs were stained for different pluripotency markers. Scale bar = 50 µm. Abbrev.: bvFTD = behavioral variant frontotemporal dementia; HC = healthy control; HRE = hexanucleotide repeat expansion; iPSCs = induced pluripotent stem cells; NANOG = Nanog homeobox; POU5F1 = POU domain, class 5, transcription factor 1; SSEA4 = Stage-specific embryonic antigen 4; TRA181 = Tumor-related Antigen-1-81

**Supplementary Fig. 3. Quality control of iPSCs from healthy controls, sporadic (C9-) and *C9orf72* HRE-carrying bvFTD (C9+) patients.** (a) A representative image of iPSCs from which the iMG were generated in this study. iMG were incubated with Vybrant dye (nuclear stain; green). Scale bar = 400 µm. (b) RNA sequencing data (mean ± standard deviation of variance stabilized counts) show expression of different stem cell marker genes in iPSCs (blue) but not in the differentiated iMG (red) for each donor. Data on iMG RNA are shown from one differentiation batch. The individual iMG lines are indicated by different symbols at the bottom. Abbrev.: bvFTD = behavioral variant frontotemporal dementia; HC = healthy control; HRE = hexanucleotide repeat expansion; iMG = induced pluripotent stem cell-derived microglia; iPSCs = induced pluripotent stem cells; *NANOG* = Nanog homeobox; *POU5F1* = POU domain, class 5, transcription factor 1; *SOX2* = SRY (sex determining region Y)-box 2

**Supplementary Fig. 4. Karyotypes of iPSCs from healthy controls, sporadic (C9-) and *C9orf72* HRE-carrying bvFTD (C9+) patients.** The karyotypes from all lines are shown. Abbrev.: bvFTD = behavioral variant frontotemporal dementia; HC = healthy control; HRE = hexanucleotide repeat expansion; iPSCs = induced pluripotent stem cells

**Supplementary Fig. 5. Timeline for iMG differentiation.** Days of the differentiation protocol are indicated by the circles. Days of freezing and thawing of erythromyeloid progenitors and subsequent experiments with iMG are indicated in black. Experimental days with cells, which were not frozen and thawn in between are shown with bright gray lines. Abbrev.: FISH = fluorescence *in situ* hybridization; IF = immunofluorescence; iMG = induced pluripotent stem cell-derived microglia; WB = Western blot

**Supplementary Fig. 6. TDP-43 translocation analysis in iMG.** (a) Representative microscopy images of TDP-43 translocation types are shown. Yellow outlines depict areas of DAPI-positive nuclei and phalloidin-positive areas, indicating cell boundaries. Scale bar = 5 µm. For statistical testing, Kruskal-Wallis test followed by Dunn’s multiple comparisons test (b; *H* = 170.7, p < 0.0001, n [type 1] = 9, n [type 2] = 391, n [type 3] = 70, n [type 4] = 3) and two-tailed Mann-Whitney test (c; test statistics = 712, n [non-pathological] = 400, n [pathological] = 73) were used. (d) Similar ratios of non-pathological *vs.* pathological TDP-43 translocation status were detected in healthy control iMG (98.539% vs. 1.461%), sporadic bvFTD iMG (98.277% vs. 1.723%), and *C9orf72* HRE-carrying bvFTD iMG (98.759% vs. 1.241%), respectively, without significantly different distribution (p = 0.3630, χ^2^ = 2.027, degree of freedom = 2). Data are shown from three independent differentiation batches. The individual iMG lines are indicated by different symbols at the bottom. Abbrev.: DAPI = 4′,6-diamidino-2-phenylindole; iMG = induced pluripotent stem cell-derived microglia; TDP-43 = TAR DNA-binding protein 43

**Supplementary Fig. 7. Principal component analysis of normalized RNA sequencing data.** Principal component analysis revealed variation between sequencing batches to be corrected for in the differential expression analysis. Abbrev.: PC = principal component

**Supplementary Fig. 8. Myeloid/microglial markers in iMG from healthy controls, sporadic (C9-) and *C9orf72* HRE-carrying (C9+) bvFTD patients.** (a) iMG were stained for myeloid lineage/microglial marker proteins. (b) Negative control staining without primary antibodies (secondary antibody only). Scale bar = 25 µm. Data are shown from one differentiation batch. Abbrev.: bvFTD = behavioral variant frontotemporal dementia; CX3CR1 = CX3C chemokine receptor 1; HC = healthy control; HRE = hexanucleotide repeat expansion; IBA1 = ionized calcium binding adapter molecule 1; iMG = induced pluripotent stem cell-derived microglia; iPSC = induced pluripotent stem cell; P2RY12 = P2Y purinoceptor 12; TMEM119 = transmembrane protein 119; TREM2 = Triggering receptor expressed on myeloid cells 2

**Supplementary Fig. 9. Assessment of RNA foci in the iMG of *C9orf72* HRE-carrying (C9+) bvFTD patients.** Microscopy images showing intranuclear RNA foci, detected by fluorescence *in situ* hybridization (FISH) using the TYE^TM^ 563(CCCCGG)_3_ probe (red) and indicated by the blue arrow. Nuclei were counterstained with DAPI (blue). Data are shown from one differentiation batch. Scale bar = 5 µm. Abbrev.: bvFTD = behavioral variant frontotemporal dementia; DAPI = 4′,6-diamidino-2-phenylindole; HRE = hexanucleotide repeat expansion; iMG = induced pluripotent stem cell-derived microglia

**Supplementary Fig. 10. TDP-43 Western Blots from iMG of healthy controls, sporadic (C9-) and *C9orf72* HRE-carrying (C9+) bvFTD patients.** (a-c) Uncropped Western blot images (showing total TDP-43 signals) are shown from three independent differentiation batches for individual iMG lines. Signal was enhanced after image acquisition to check for faint signals emerging from potential C-terminal fragments. The presence of C-terminal TDP-43 fragments (∼35 and ∼25 kDa) was not evident on the blots. Abbrev.: bvFTD = behavioral variant frontotemporal dementia; HC = healthy control; HRE = hexanucleotide repeat expansion; iMG = induced pluripotent stem cell-derived microglia; kDa = kilo Dalton

**Supplementary Fig. 11. Phagocytosis-related gene expression in iMG of healthy controls, sporadic (C9-) and *C9orf72* HRE-carrying (C9+) bvFTD patients.** (a) RNA levels according to RNA sequencing are shown for each donor as mean ± standard deviation of variance stabilized counts of *CLEC7A*: *F* = 0.3732, *p* = 0.7035; *TLR2*: *F* = 0.6503, *p* = 0.5551; *TLR5*: *F* = 4.926, *p* = 0.0542; *TLR6*: *F* = 0.02323, *p* = 0.9771; *n* [HC] = 3; *n* [bvFTD C9-] = 3; *n* [bvFTD C9+] = 3; One-way ANOVA followed by Tukey’s multiple comparisons test. (b) Heatmap shows the comparison-wise fold changes (log2) for all genes in the Gene ontology biological process phagocytosis category (GO 0006909), phagolysosome (GO 0032010), autophagosome-lysosome fusion (GO:0061909), and the Reactome category autophagy (R-HSA-9612973) that are differentially expressed (FDR < 0.05) in any of the three comparisons. Significance levels: * = 0.01 ≤ padj < 0.05, ** = 0.001 ≤ padj < 0.01, *** = padj < 0.001. Data are shown from one differentiation batch. The individual iMG lines are indicated by different symbols at the bottom. Abbrev.: bvFTD = behavioral variant frontotemporal dementia; *CLEC7A* = C-type lectin domain containing 7A; HC = healthy control; HRE = hexanucleotide repeat expansion; iMG = induced pluripotent stem cell-derived microglia; *TLR* = Toll-like receptor

**Supplementary Fig. 12. Inflammation-related gene expression in iMG of healthy controls, sporadic (C9-) and *C9orf72* HRE-carrying (C9+) bvFTD patients.** (a) Heatmap shows the comparison-wise fold changes (log2) for all genes in the Gene ontology biological process acute inflammatory response (GO 0002526) that are differentially expressed (FDR < 0.05) in any of the three comparisons. (b) RNA levels of *CCL2*: *F* = 1.767, *p* = 0.2493; *CHIT1*: *F* = 1.252, *p* = 0.3511; *CXCL10*: *F* = 1.230, *p* = 0.3567; *ILB*: *F* = 7.063, *p* = 0.0265; *IL18*: *F* = 0.08600, *p* = 0.9187; *TNF*: *F* = 3.261, *p* = 0.1100. Data are shown for each donor as mean ± standard deviation of variance stabilized counts; *n* [HC] = 3; *n* [bvFTD C9-] = 3; *n* [bvFTD C9+] = 3. Significance levels: * = 0.01 ≤ padj < 0.05, ** = 0.001 ≤ padj < 0.01, *** = padj < 0.001. Data are shown from one differentiation batch. The individual iMG lines are indicated by different symbols at the bottom. Abbrev.: bvFTD = behavioral variant frontotemporal dementia; *CCL2* = C-C motif chemokine ligand 2; *CHIT1* = chitinase 1; *CXCL10* = C-X-C motif chemokine ligand 10; HC = healthy control; HRE = hexanucleotide repeat expansion; *IL* = interleukin; iMG = induced pluripotent stem cell-derived microglia: *TNF* = tumor necrosis factor

**Supplementary Video 1. Phagocytosis assay of Con_1 line without prior serum starvation (s. online)**

**Supplementary Video 2. Phagocytosis assay of Con_1 line after 24 h serum starvation (s. online)**

**Supplementary Video 3. Phagocytosis assay of Con_2 line without prior serum starvation (s. online)**

**Supplementary Video 4. Phagocytosis assay of Con_2 line after 24 h serum starvation (s. online)**

**Supplementary Video 5. Phagocytosis assay of Con_3 line without prior serum starvation (s. online)**

**Supplementary Video 6. Phagocytosis assay of Con_3 line after 24 h serum starvation (s. online)**

**Supplementary Video 7. Phagocytosis assay of Ftd_1 line without prior serum starvation (s. online)**

**Supplementary Video 8. Phagocytosis assay of Ftd_1 line after 24 h serum starvation (s. online)**

**Supplementary Video 9. Phagocytosis assay of Ftd_2 line without prior serum starvation (s. online)**

**Supplementary Video 10. Phagocytosis assay of Ftd_2 line after 24 h serum starvation (s. online)**

**Supplementary Video 11. Phagocytosis assay of Ftd_4 line without prior serum starvation (s. online)**

**Supplementary Video 12. Phagocytosis assay of Ftd_4 line after 24 h serum starvation (s. online)**

**Supplementary Video 13. Phagocytosis assay of Ftd_5 line without prior serum starvation (s. online)**

**Supplementary Video 14. Phagocytosis assay of Ftd_5 line after 24 h serum starvation (s. online)**

**Supplementary Video 15. Phagocytosis assay of Ftd_6 line without prior serum starvation (s. online)**

**Supplementary Video 16. Phagocytosis assay of Ftd_6 line after 24 h serum starvation (s. online)**

