## Supplementary Figures 1-12 for "*C9orf72* repeat expansion-carrying iPSC-microglia from FTD patients show increased phagocytic activity concomitantly with decreased number of autophagosomal-lysosomal vesicles"

Supplementary Figure 1.

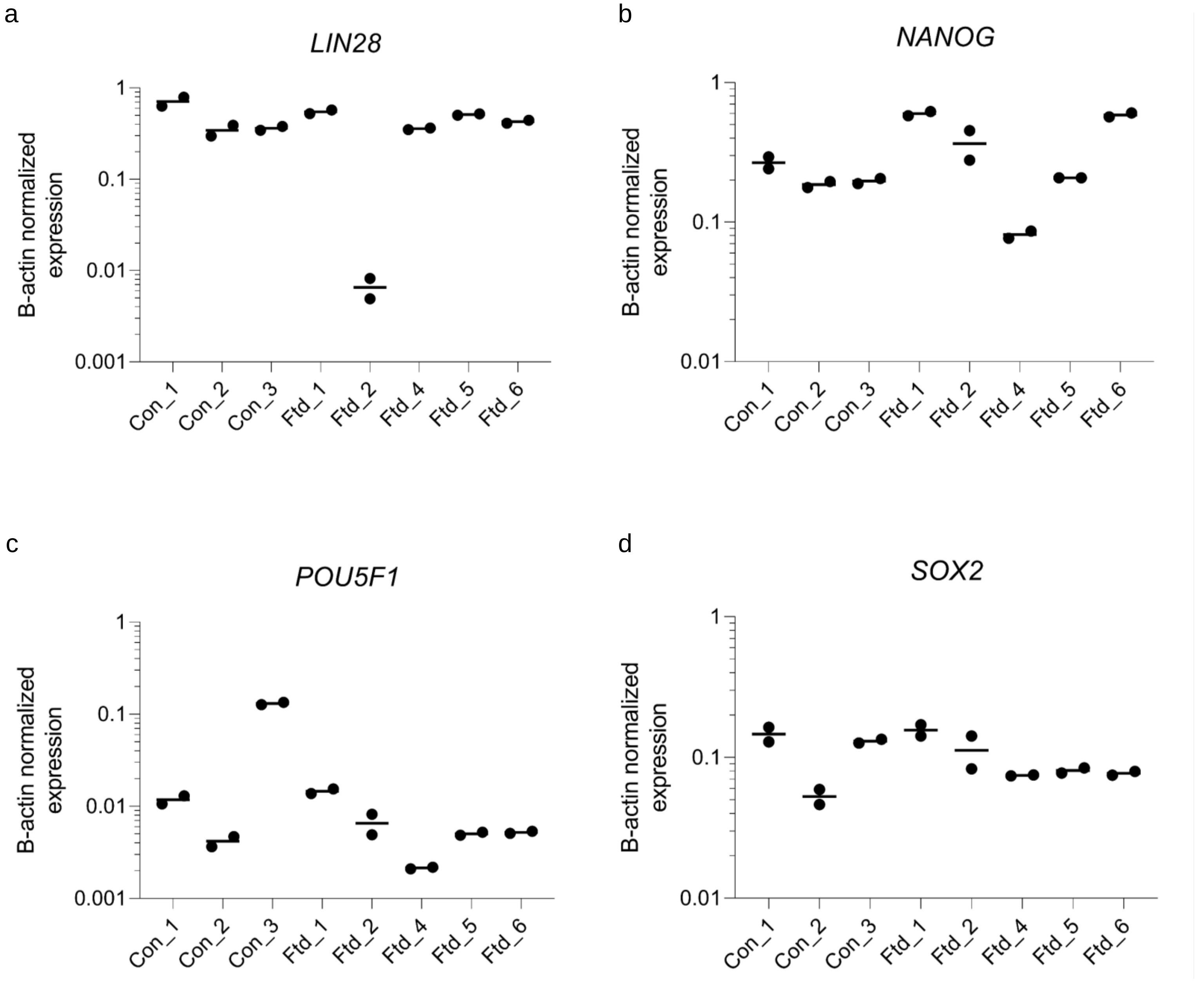

Supplementary Figure 2.

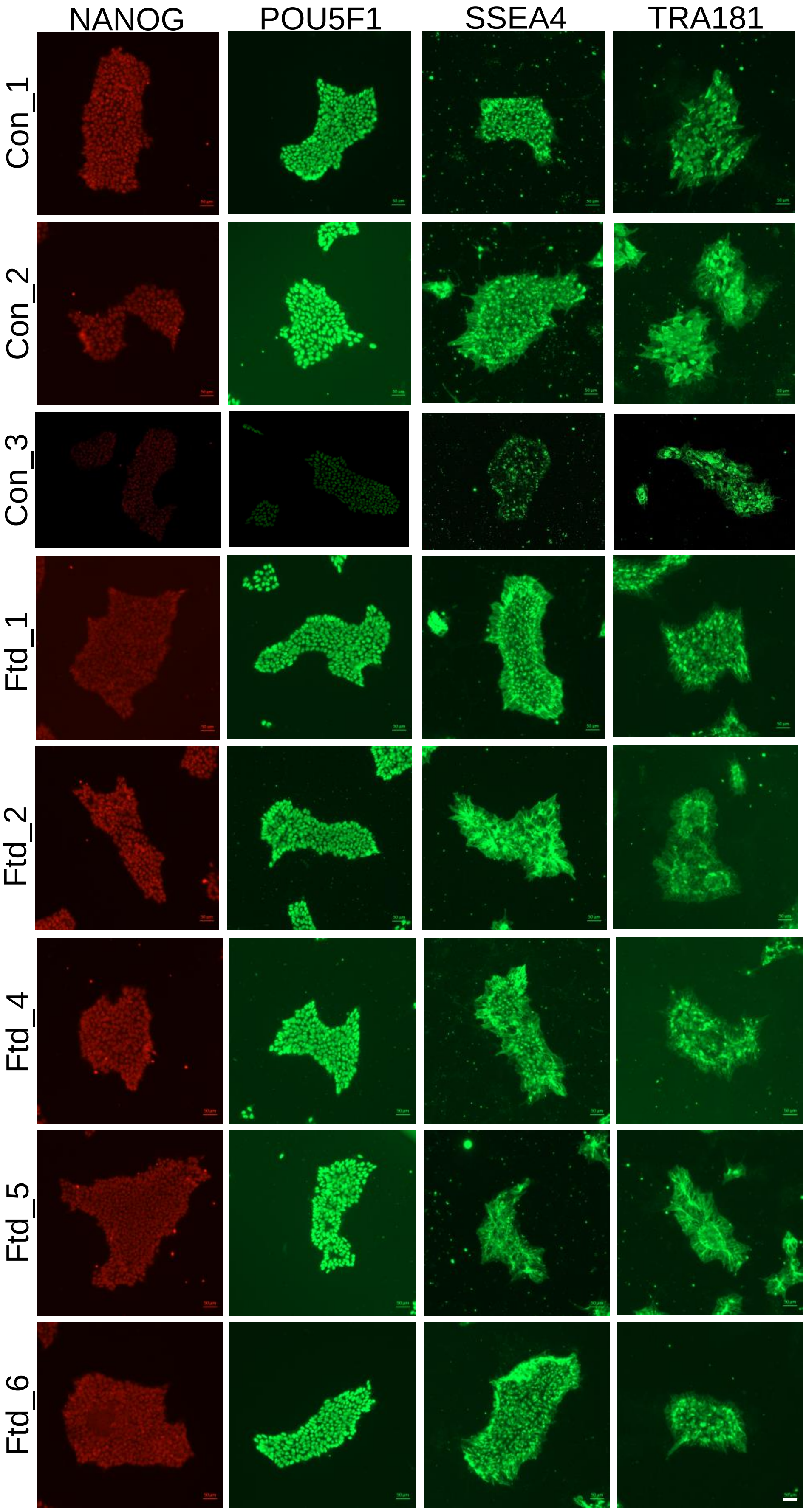

Supplementary Figure 3.

a

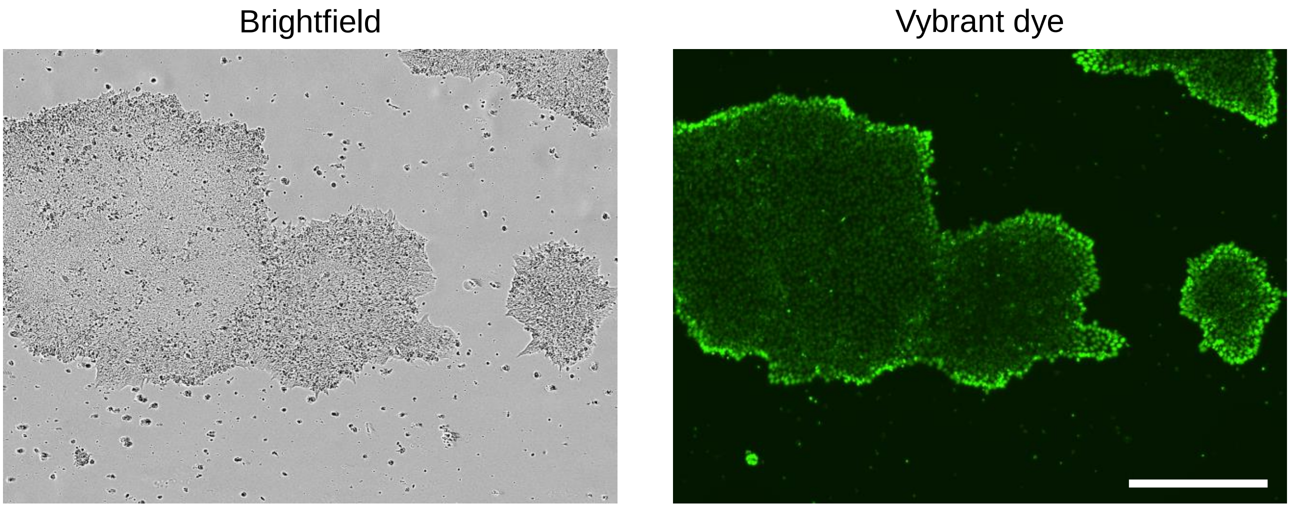


b

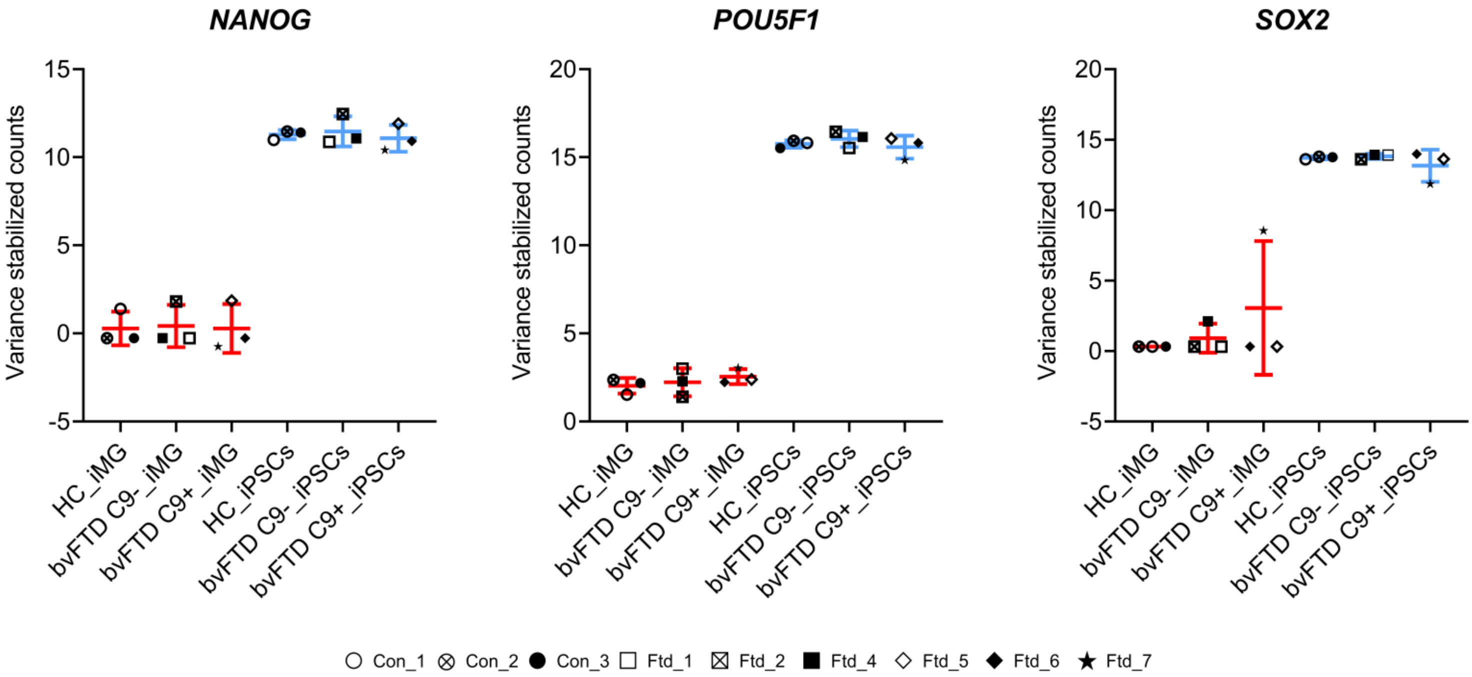

Supplementary Figure 4.

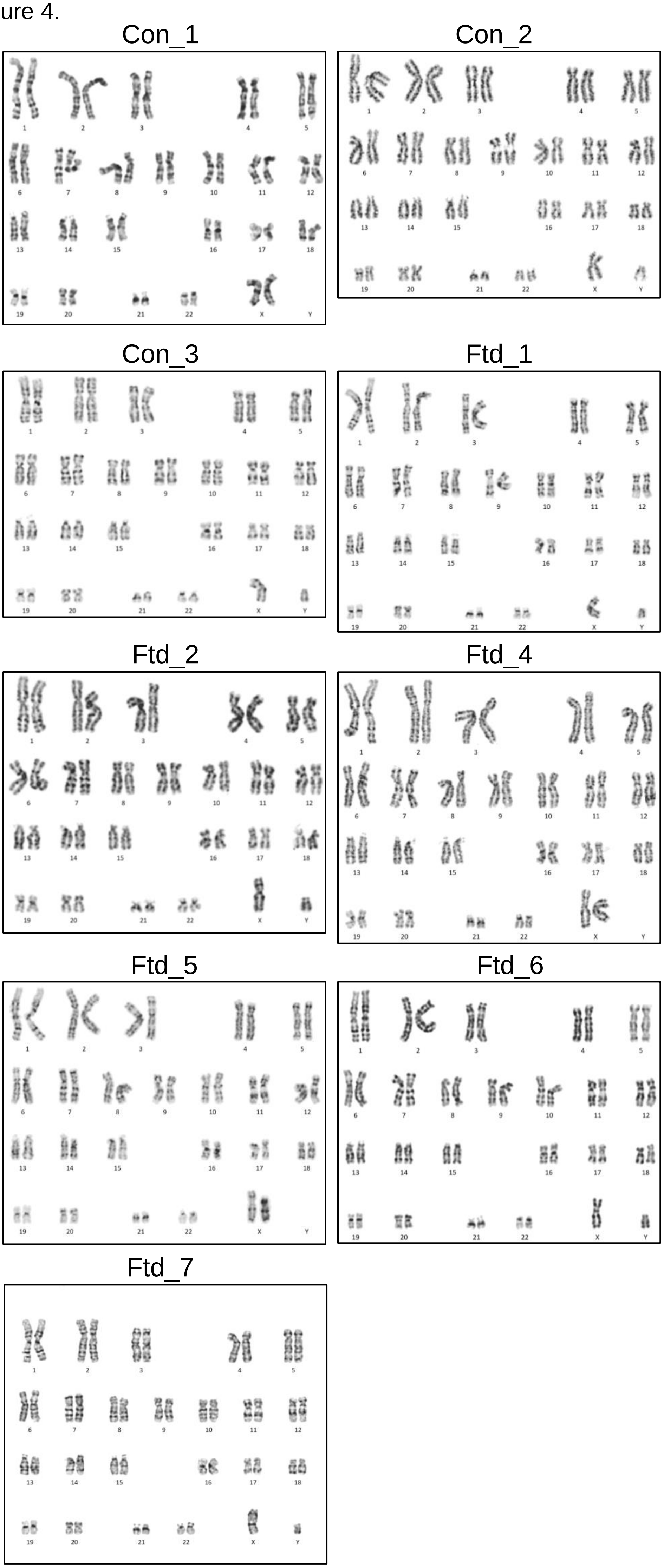

Supplementary Figure 5.

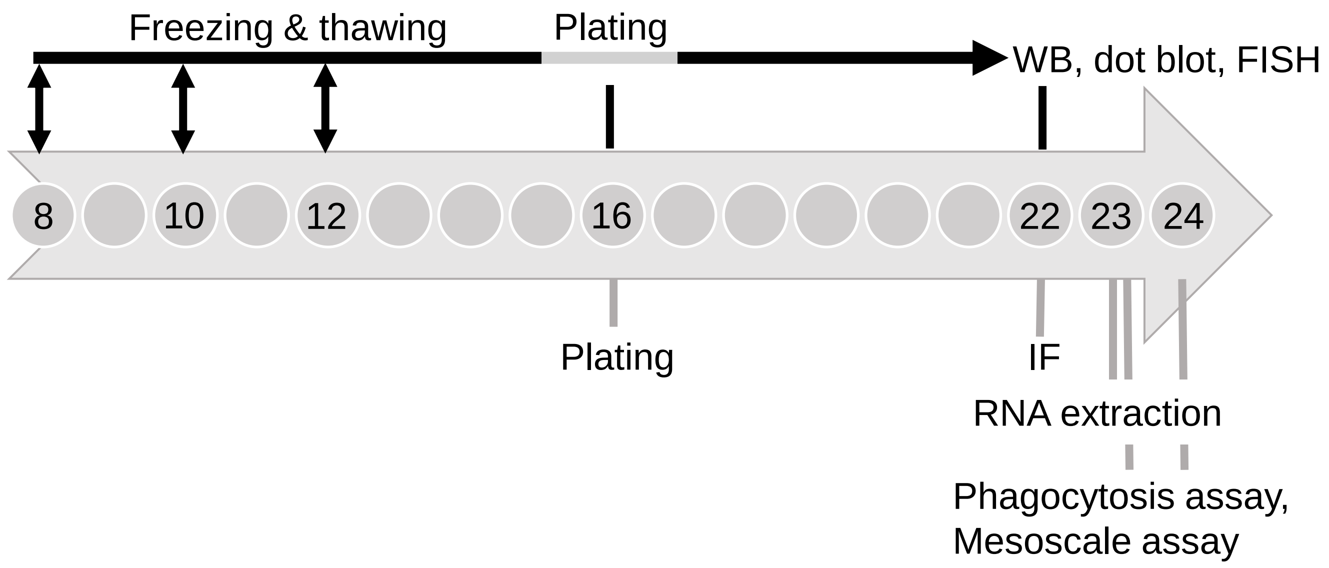

Supplementary Figure 6.

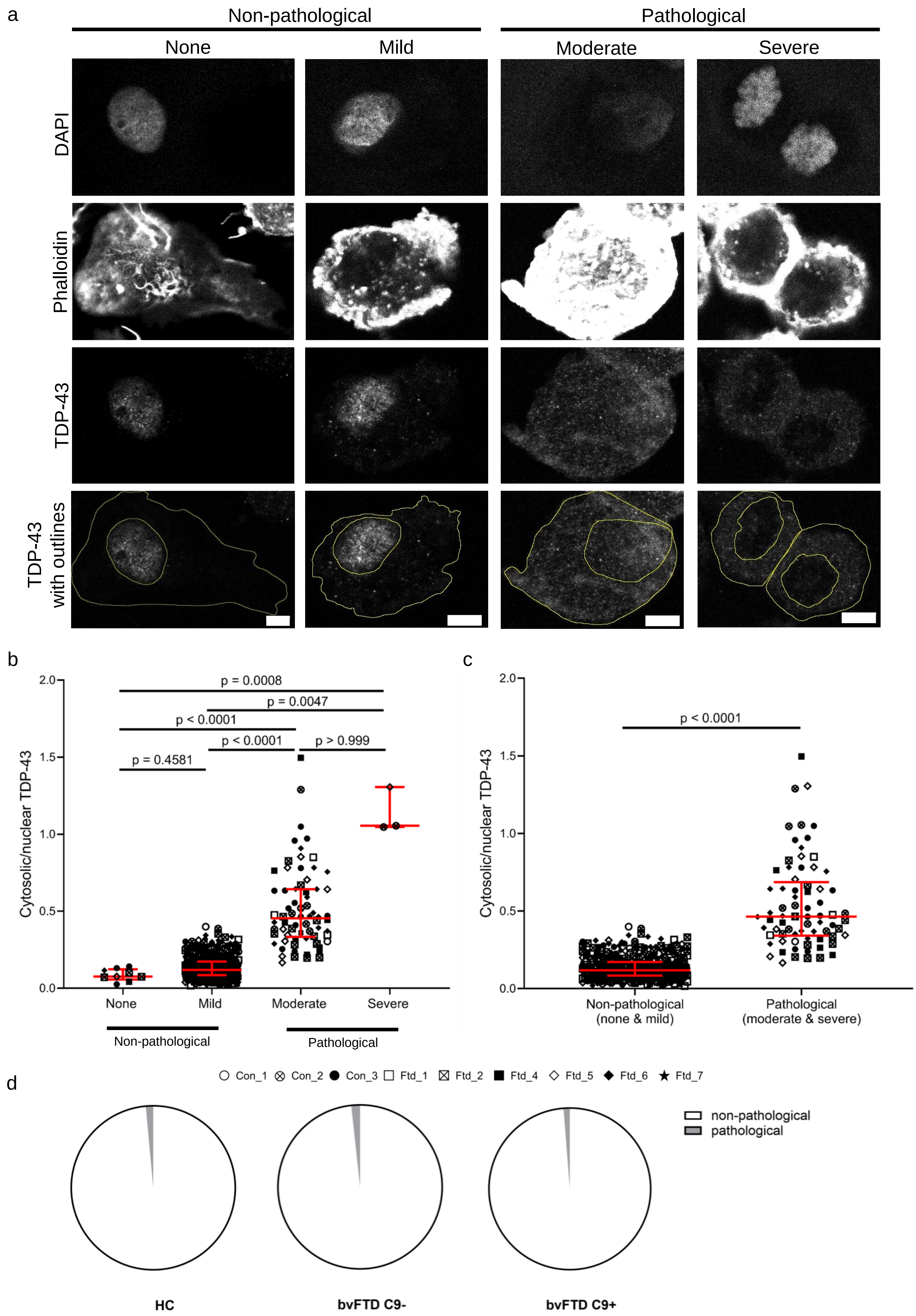

Supplementary Figure 7.

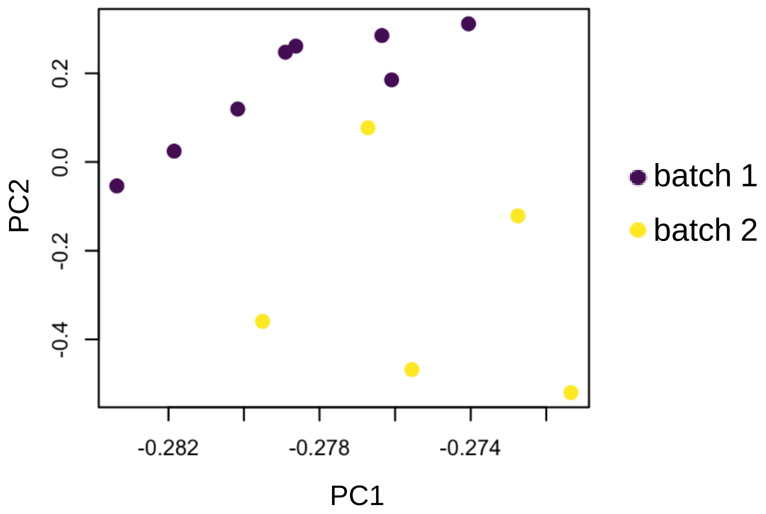

Supplementary Figure 8.

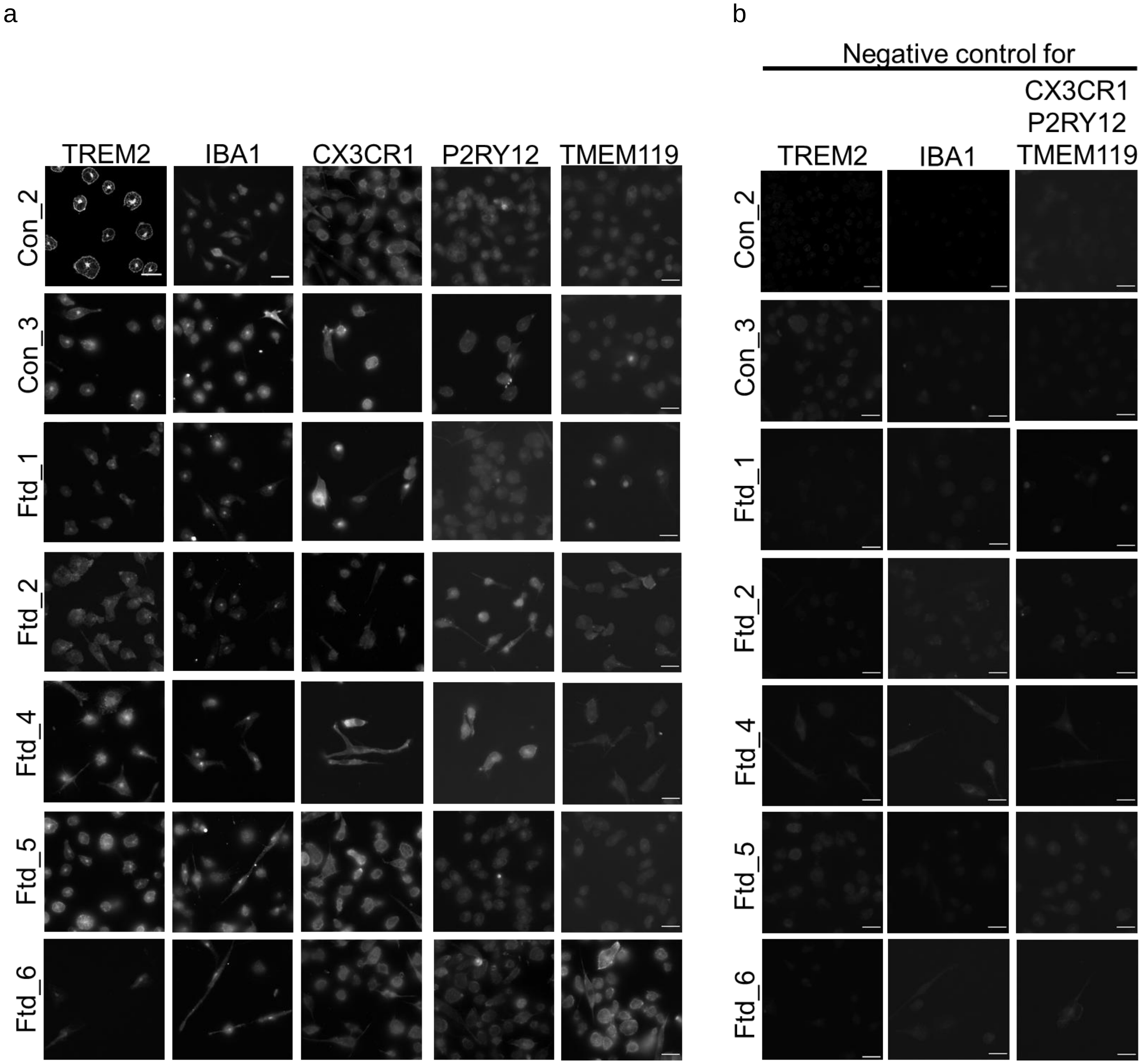

Supplementary Figure 9.

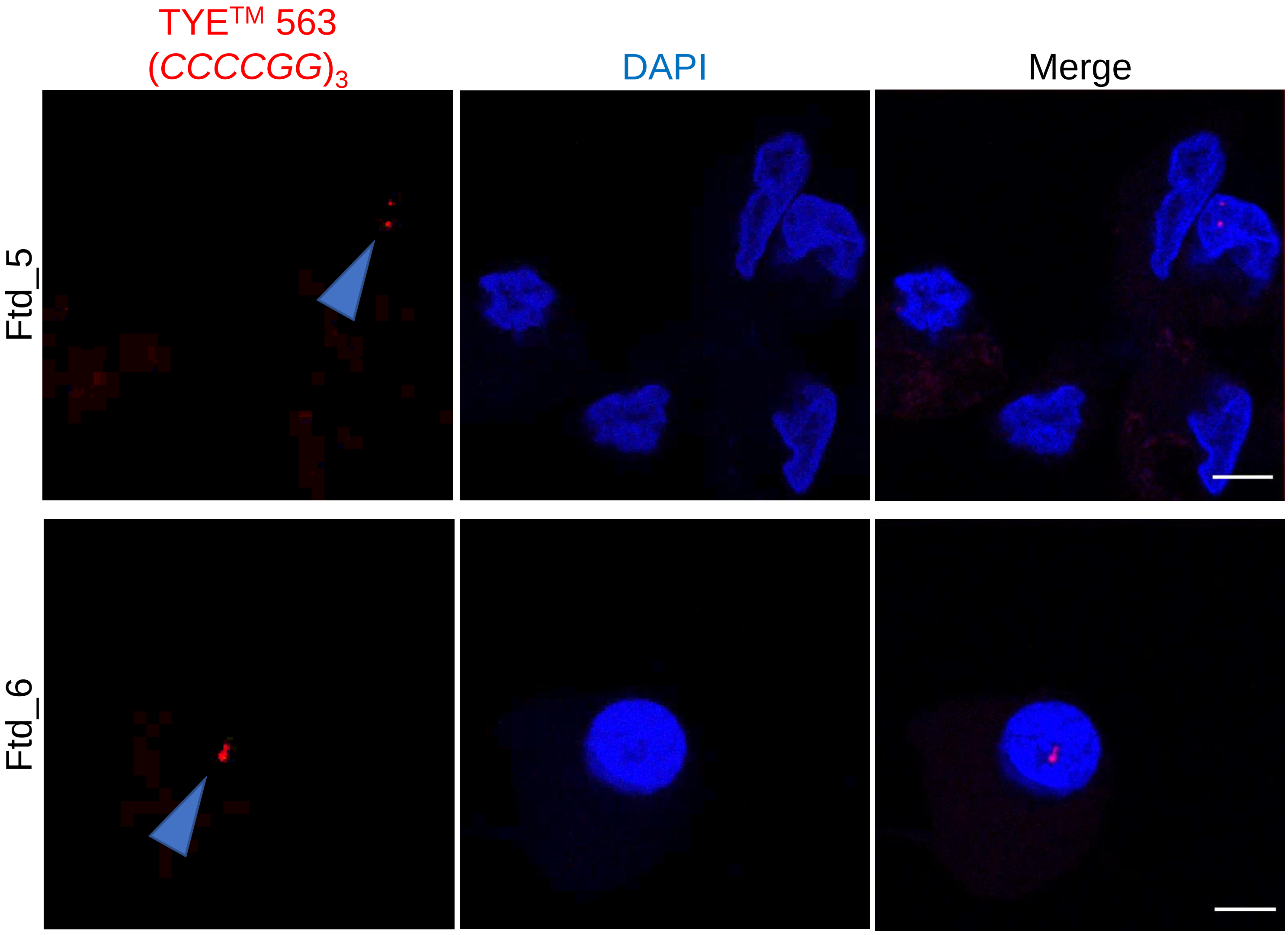

Supplementary Figure 10.

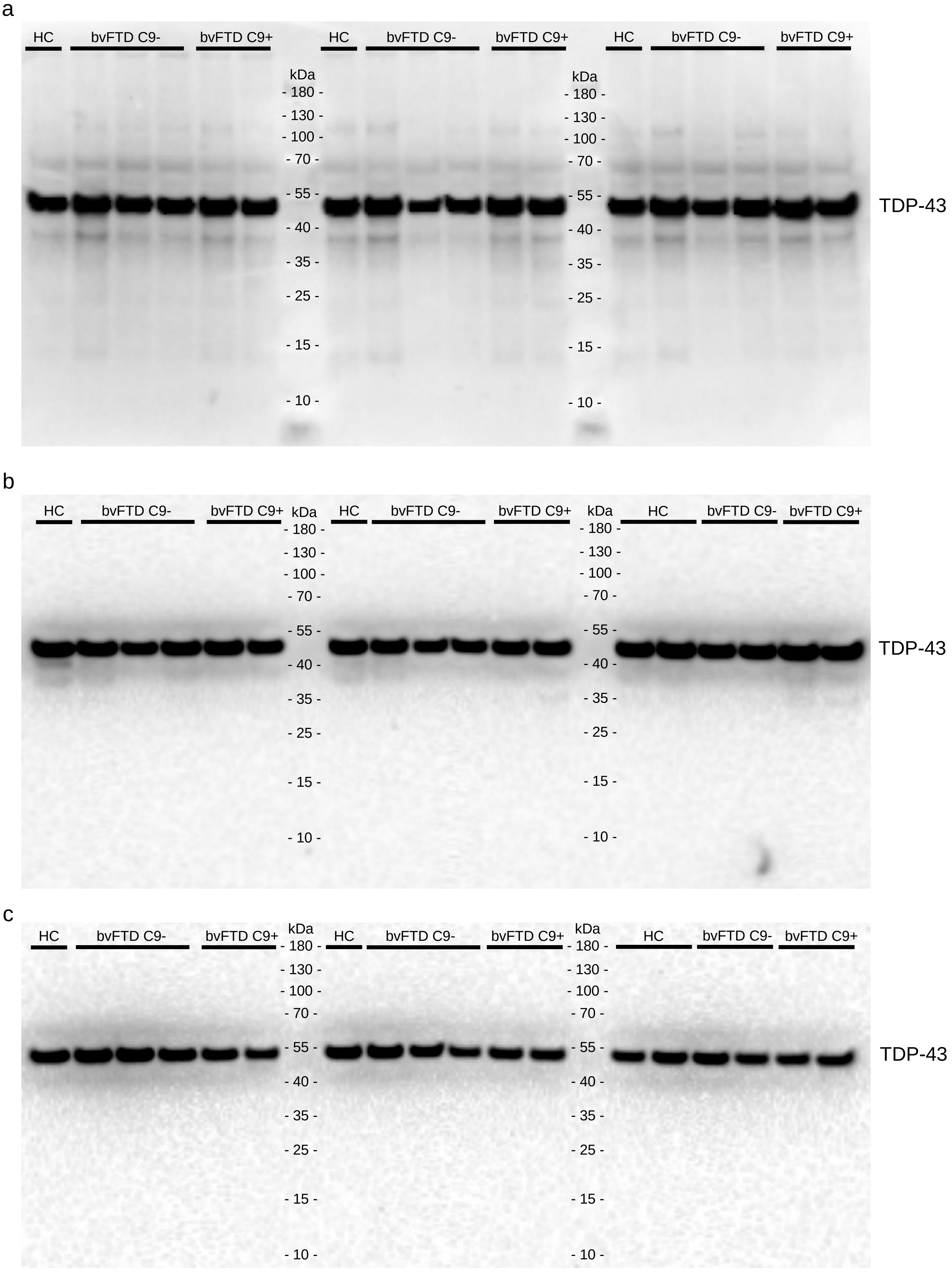

Supplementary Figure 11.

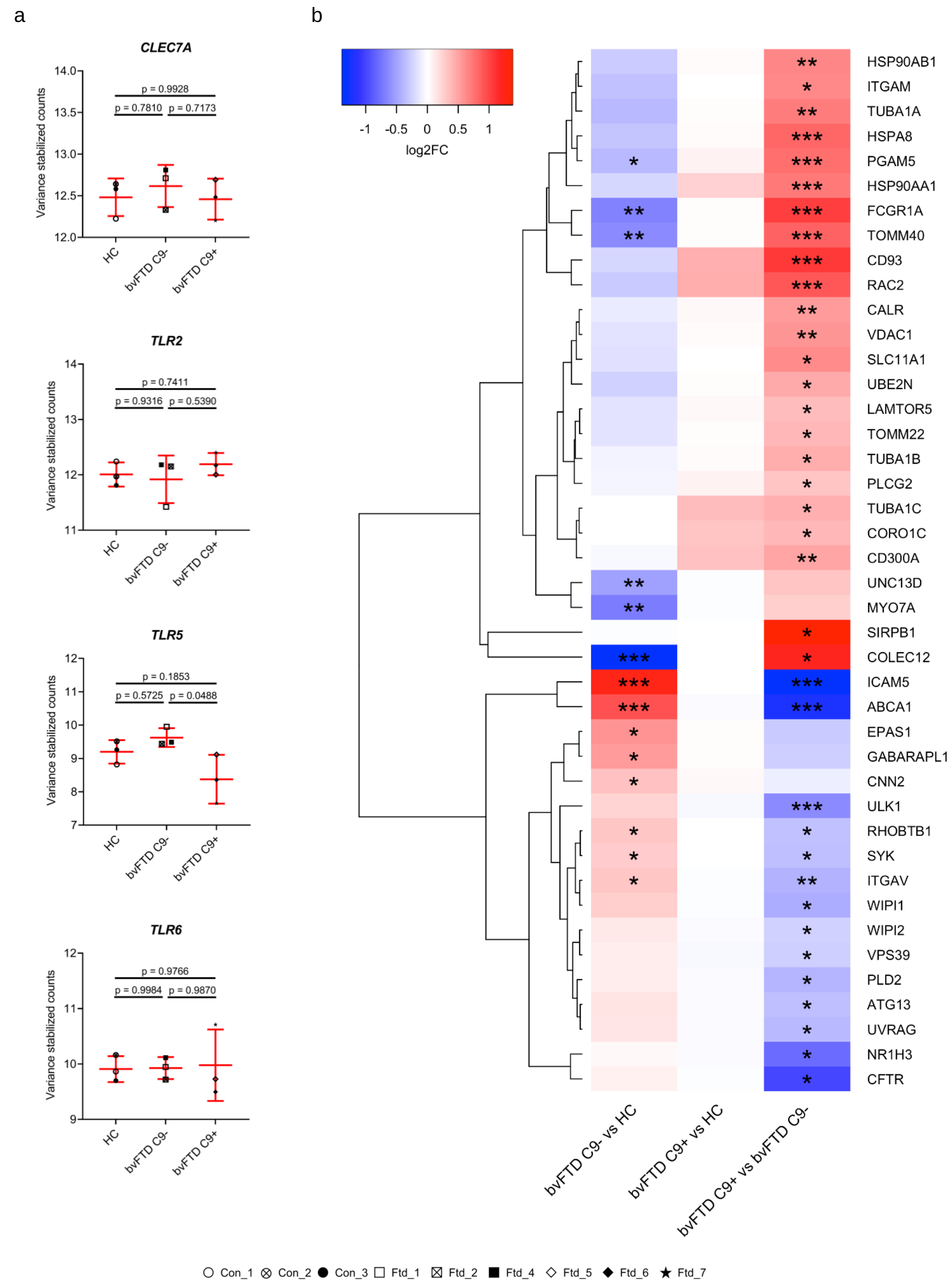

Supplementary Figure 12.

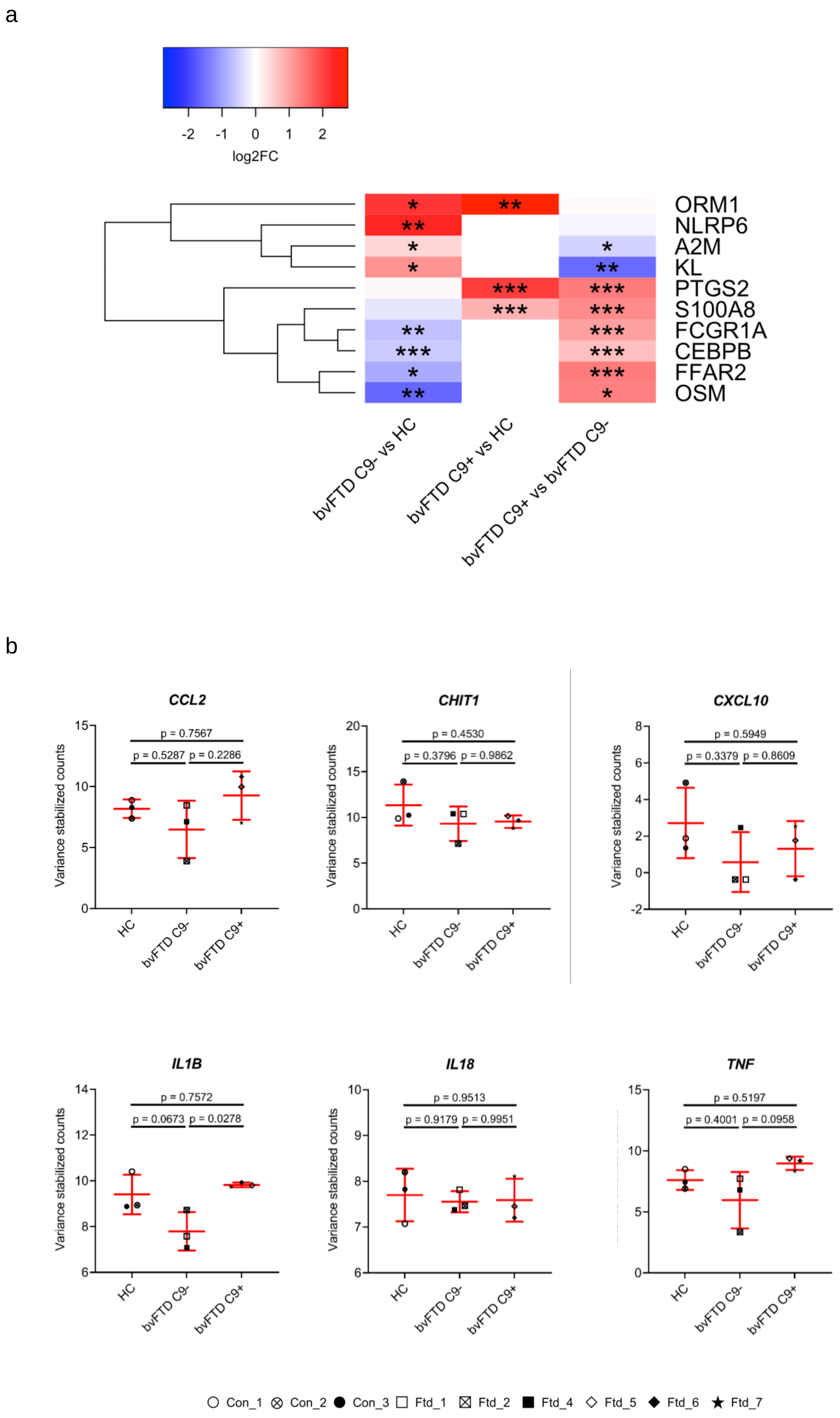
